# “A decrease in specific health-associated commensals is linked to progressive periodontal tissue destruction independent of dysbiotic community profiles”

**DOI:** 10.64898/2026.04.30.721960

**Authors:** Natalia Endo, Joaquin Espinoza-Arrue, Marion Arce, Nicole Traver, Maria I. Muñoz-Sepúlveda, Daniel Sansores-España, Valentina Olmedo, Catalina Moreno, Javiera Canelo, Montserrat Reyes, Alex M Valm, Nicolas Dutzan, Loreto Abusleme

## Abstract

Periodontitis is a chronic inflammatory disease associated with dysbiotic microbial communities that leads to destruction of the tooth-supporting tissues. The transition from host-microbial periodontal homeostasis to disease remains poorly understood. The murine ligature-induced periodontitis model was employed to characterize the temporal dynamics of the subgingival microbiome and host tissue features. Ligatures were placed in C57BL/6N mice, and collected on days 0, 1, 3, 5, and 7 post-induction. Bacterial load, alveolar bone loss, immune cells (CD45⁺), cells with osteoclastogenic potential (TRAP⁺) and collagen destruction were analyzed. Additionally, the V4 region of the 16S rRNA gene was sequenced for ecological analyses, including co-occurrence networks and functional prediction. Spatial distribution of the most abundant species was visualized using CLASI-FISH microscopy. Finally, association models were performed to link bacterial abundances with time and tissue parameters. The most substantial microbial shift occurred on day 1, and a dysbiotic community was established by day 3. CD45⁺ cell infiltration increased as early as day 1, preceding the rise in TRAP⁺ cells on day 3 and the onset of tissue destruction on day 5. By day 7, predicted bacterial functions included protein export, lipid and galactose metabolism. Health-associated taxa were identified, and their abundance correlated positively with collagen integrity and negatively with immune cell infiltration and bacterial load, highlighting their role in homeostasis. These findings provide a high-resolution temporal map of microbiome-host interactions during experimental periodontitis establishment and identify specific microbial and cellular windows for potential therapeutic intervention.

## Introduction

Periodontitis is a highly prevalent inflammatory disease characterized by progressive destruction of tooth-supporting tissues, including alveolar bone and oral mucosa, being the leading cause of tooth loss among adults and a contributor to adverse systemic health outcomes ^1–4^. In this condition, the establishment of a dysbiotic biofilm triggers an inflammatory response that drives the activation and recruitment of immune cells ^5,6^. This immune-destructive process is complex, involving changes in the oral microbial community alongside immune and histopathological events that collectively shape disease development ^7,8^. Accordingly, the transition from health to disease involves a shift in the microbiome community structure, characterized not only by increased representation of pathobionts but also by changes in the functional properties and pathogenic potential of the microbial community ^5,9,10^. In parallel, the inflammatory response is coordinated by a multicellular network in which immune cells interact with structural cells, forming spatially organized niches that define site-specific immune characteristics at the oral mucosa ^11,12^.

While pre-clinical and clinical studies have provided a detailed cross-sectional view of disease-associated features, they are inherently limited in capturing the dynamic processes underlying disease progression. Given these limitations, longitudinal and experimental models are essential tools for elucidating temporal and mechanistic events driving the establishment of periodontitis. In particular, animal models of periodontitis enable the characterization of underlying processes that remain difficult to assess in human clinical research ^13^, thereby providing critical insights into disease pathogenesis.

The ligature-induced periodontitis (LIP) model in mice is one of the most employed to study periodontitis and aims to reproduce human periodontal pathology, defined by an increased bacterial load in the ligature and a dysbiotic state associated with progressive periodontal tissue destruction ^6,14–16^. This model allows for the study of host-microbiome interactions, in which shifts in microbial communities are functionally linked to immune responses that drive inflammatory bone loss ^6^. This change in the microbiome is characterized by a dysbiotic state with reduced alpha diversity, which also impacts beta diversity. Such dysbiosis differs in microbial composition across studies but remains consistently associated with alveolar bone loss, with *Streptococcus danieliae* characterizing periodontal health and taxa such as *Enterococcus faecalis* and *Faecalibaculum rodentium* predominating in disease ^15^.

Consistently, the immune response observed in the LIP model resembles that of periodontitis, where dysbiotic communities drive inflammatory bone loss through Th17 responses, a pathway whose impairment in humans is associated with reduced disease severity, supporting its conserved function across species ^6^.

Given its relevance, the LIP model has been used in longitudinal studies to evaluate microbial changes in the ligature and immune-inflammatory responses over time ^17–23^. However, most of these studies have relied on ecological microbiome analyses with limited taxonomic resolution and have primarily examined microbial and host responses independently. Therefore, an in-depth understanding of microbial dynamics and their relationship with tissue alterations during disease establishment remains to be explored.

In this work, the LIP model was employed to elucidate the development of experimental periodontitis through a longitudinal manner that combines detailed characterization of the microbiome with histological changes. By integrating community ecology, functional potential, host–microbe associations across time, and spatial organization assessed by CLASI-FISH, this analytical framework enables the identification of ecological shifts and their temporal association with immune cell infiltration and tissue damage. This approach provides a more comprehensive understanding of the critical events underlying disease establishment, further illuminating potential areas for prevention and intervention.

## Results

### An increase in total bacterial load and inflammatory cell infiltration precedes tissue destruction during LIP

Total bacterial load, alveolar bone loss, and histological changes were assessed during LIP on days 0 (D0), 1 (D1), 3 (D3), 5 (D5), and 7 (D7) post-ligation. A significant increase in bacterial load was observed on days 3, 5, and 7 compared to day 0 (2 hours of ligature placement) (p=0.0025; <0.0001; 0.0017) (Fig. 1A). This increase preceded the detection of significant alveolar bone loss, which was observed on D5 and D7 during LIP (p=0.0010; <0.001) (Fig. 1B, S1A). Furthermore, a progressive accumulation of immune cells in periodontal tissues was observed (Fig. S1B), as reflected by a significant increase in the percentage of immune (CD45^+^) cells at D1, D3, D5, and D7 compared to D0 (p=0.0261; 0.0008; < 0.0001; < 0.0001; respectively) (Fig. 1C). The assessment of TRAP^+^ cells, a commonly used marker of osteoclast lineage cells and osteoclastogenic potential, revealed an increase in their number on D3, D5, and D7 compared to baseline (p=0.0043; <0.001; 0.0243) (Fig. 1D). To evaluate the destruction of periodontal connective tissue, we assessed collagen loss from the proximal area between adjacent teeth. We observed a decrease in collagen percentage from D5 and D7, with a gradual decline compared to baseline (p=0.0037; 0.0002) (Fig. 1E). Additionally, Brown and Brenn staining showed a progressive accumulation of bacteria in the ligature over time, without bacterial presence in the surrounding periodontal tissues (Fig. S1C).

**Figure 1.**
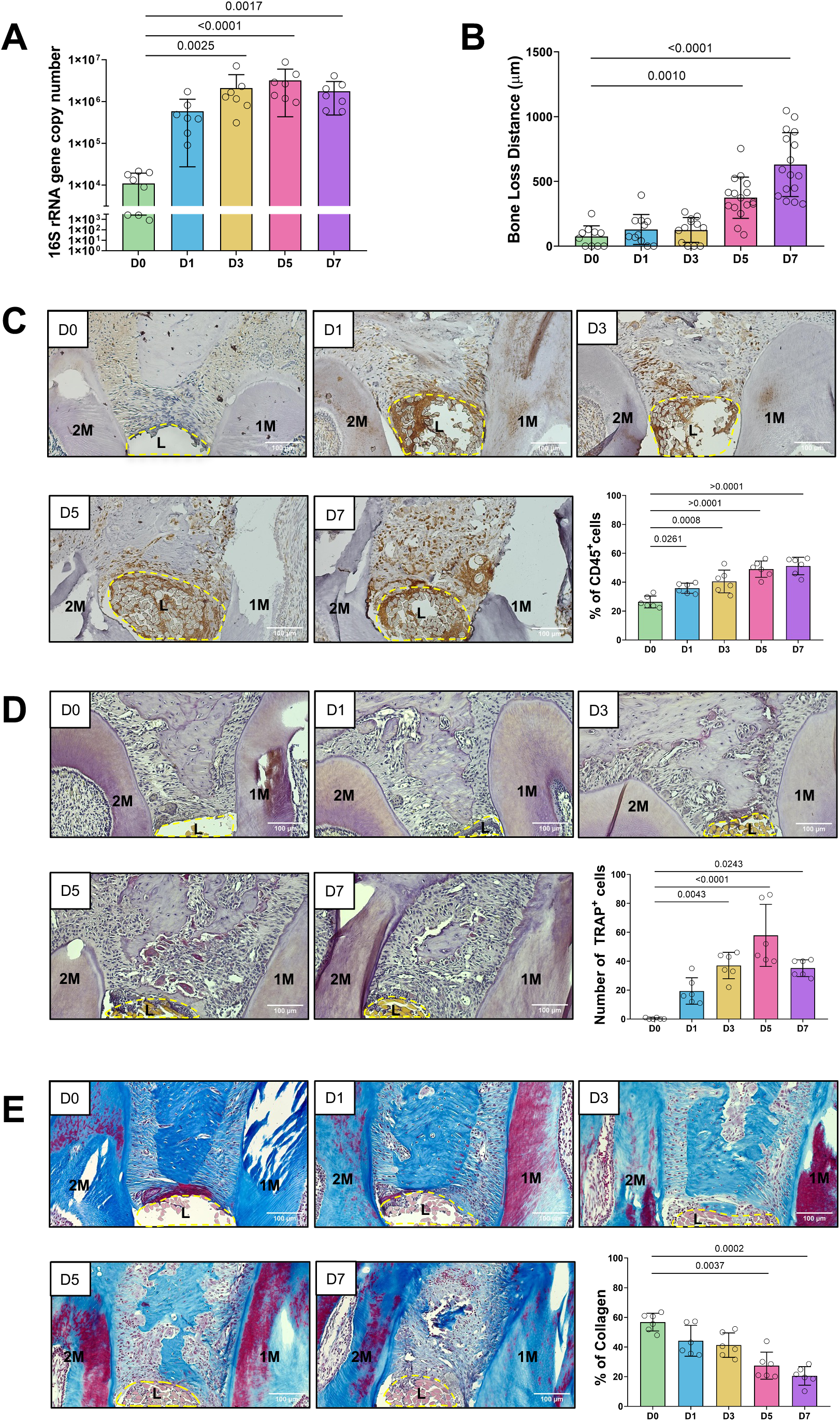
Longitudinal assessment of total bacterial load, alveolar bone loss, and periodontal tissue remodeling in ligature-induced periodontitis. (A) The bar graph depicts total bacterial load (16S rDNA gene copies per time point). (B) The bar graph depicts alveolar bone loss (in millimeters) for each time point. (C) Representative histological sections stained with immunohistochemistry, with the corresponding graph showing the percentage of CD45^+^ cells. (D) Representative histological sections stained for tartrate-resistant acid phosphatase (TRAP) to identify cells with osteoclastogenic potential, with the corresponding graph quantifying the number of TRAP^+^ cells. (E) Representative histological sections stained with Masson’s trichrome, highlighting collagen fibers in blue, with the corresponding graph showing the percentage of collagen per tissue area. For all graphs, data are presented as mean ± SEM. Statistical significance was calculated for all time-points versus the Day 0 control group, was determined using the Kruskal-Wallis test.

### The subgingival microbiome exhibits shifts in ecological dynamics within three days of periodontitis induction

No differences in microbial richness (number of ASVs) were detected between experimental timepoints (Fig. 2A). However, indices sensitive to species abundance and evenness showed a progressive decline over time, as lower diversity index values were indicated by the non-parametric Shannon index, when comparing D0-D5, D1-D3, D1-D7, and D3-D5 during LIP (Fig. 2B). Similar results were obtained using the inverse Simpson diversity index (Fig. S2A), indicating that diversity decreased over time, driven by shifts in species abundance rather than loss of taxa.

**Figure 2.**
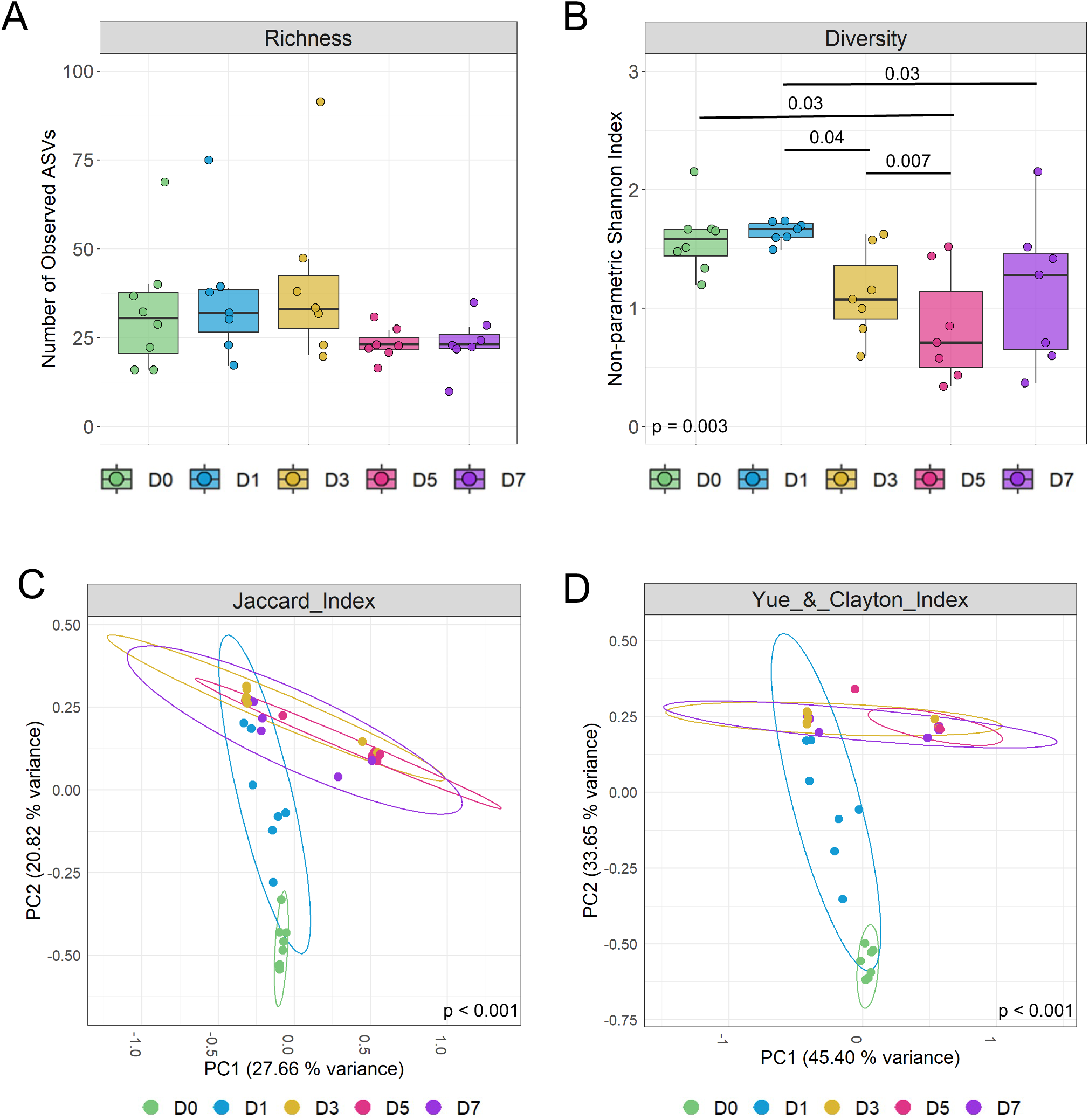
Temporal shifts in microbial alpha and beta diversity during ligature-induced periodontitis.(A) Non-parametric Shannon diversity index, a measure of community richness and evenness. Statistical significance between time points was assessed using the Kruskal-Wallis test followed by Dunn’s multiple-comparison post-hoc test. (B) Observed number of Amplicon Sequence Variants (ASVs), a measure of community richness. Statistical significance between time points was assessed using the Kruskal-Wallis test followed by Dunn’s multiple-comparisons post hoc test. (C) Principal coordinate analysis (PCoA) plot based on the Jaccard index, a qualitative measure of community composition (presence/absence). Data clouds are shown with 95% confidence ellipses. The significance of the separation among groups (p < 0.001) was determined by permutational analysis of variance (PERMANOVA). (D) Principal coordinate analysis (PCoA) plot based on the Yue & Clayton (ThetaYC) distance, a quantitative measure of community structure (abundance-weighted). Data clouds are shown with 95% confidence ellipses. The significance of the separation among groups (p < 0.001) was determined by permutational analysis of variance (PERMANOVA).

A Principal Coordinate Analysis (PCoA) was conducted to evaluate differences in microbial community structure. Significant clustering of communities by experimental time-point was observed, indicating distinct changes in microbiome structure as the transition from health to disease occurred (Fig. 2C, 2D, S2B-C). This finding was consistent across three indexes: the Jaccard index (community composition) (Fig. 2C), the Yue & Clayton measure (Fig. 2D), and the Bray-Curtis index (Fig. S2B) (both assess community structure). Additionally, the phylogenetic measures unweighted UniFrac (presence/absence of taxa) and the weighted UniFrac (community structure), clustered the communities into three distinct phylogenetic groups: baseline (D0 alone), an intermediate cluster comprising D1 and D3, and a later-stage cluster comprising D5 and D7 (Fig. S2C-D), also pointing towards the microbial transition as inflammation and tissue destruction ensue.

### The ligature-associated microbiome undergoes dysbiosis and spatial re-structuring within three days post-induction

Taxonomic profiling and linear discriminant analysis (LEfSe) revealed the bacterial dynamics underlying ecological shifts during the transition from health to disease in LIP (Fig. 3A, Fig. S3, S4). At baseline (D0), the microbial profile was characterized by a specific set of taxa consistently enriched in this state. Among these, *Achromobacter_sp.(A._xylosoxidans)_ASV3, Streptococcus_sp._(S._danieliae)_ASV4/7, Burkholderia_sp._*(*B._cenocepacia*)_*ASV10, Serratia_sp._*(*S._marcescens*)_*ASV9, and Pseudomonas_sp.(P._protegens)_ASV8* were recurrently identified as discriminant features across baseline comparisons (Fig. 3A, S3A, S4). Given their consistent association with steady state, these taxa were considered health-associated taxa (HAT).

**Figure 3.**
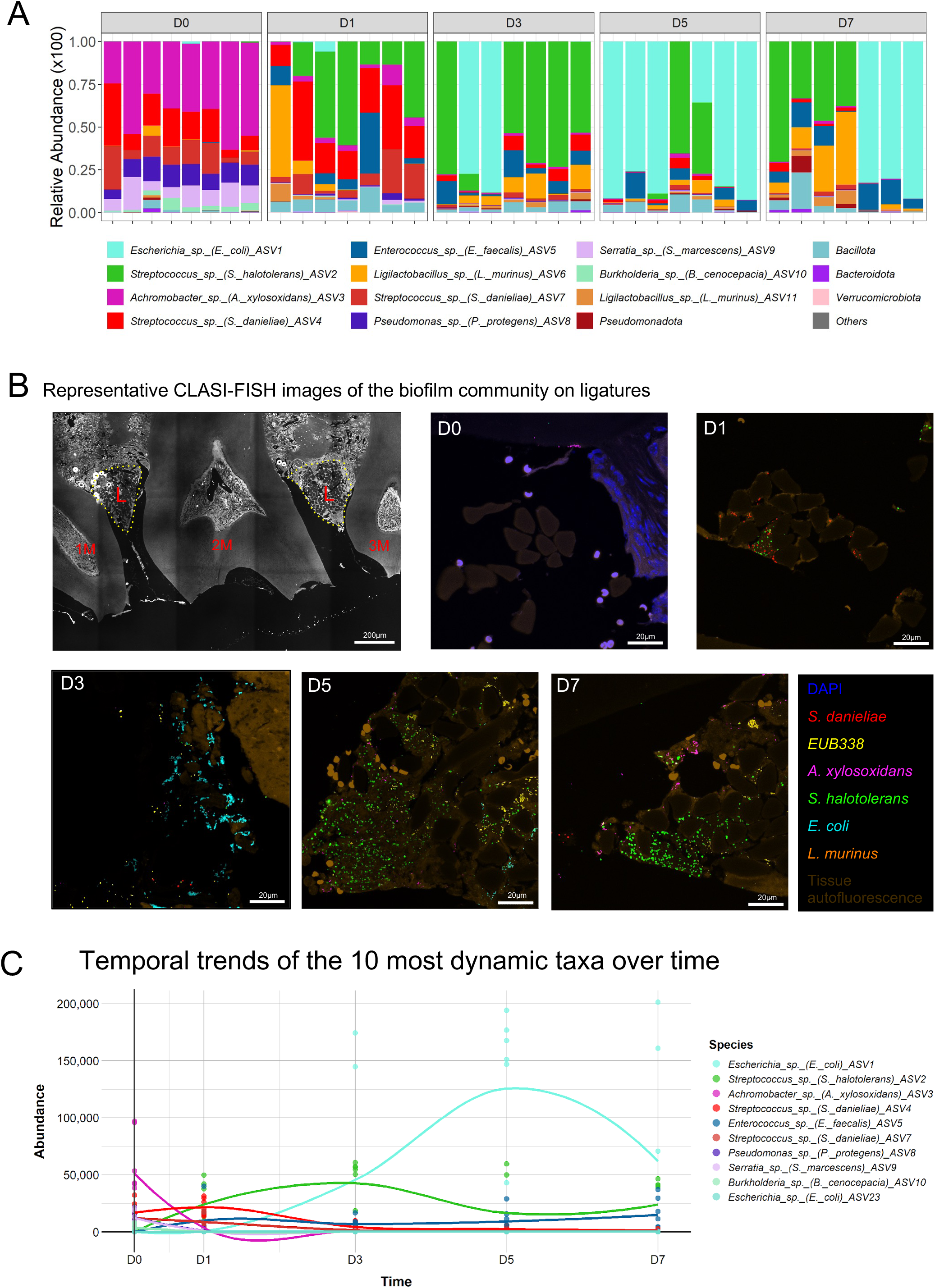
Microbial community composition and dynamics across the timeline of ligature-induced periodontitis. (A) Relative abundance of bacterial taxa at ASV level across different time points. Each bar represents an individual sample. (B) Representative CLASI-FISH images of oral bacteria on ligatures at day 0, 1, 3, 5 and 7 post-induction. Scale bar: 20 µm. Each color represents one of the most abundant ASVs. Probes target highly abundant ASVs. (C) Temporal trends of the 10 most dynamically significant taxa over time, identified by a MaAsLin 3 longitudinal analysis. The analysis was run using a Linear Model (LM) on TSS-normalized and LOG-transformed data. The reference time point was set at Day 0. Only taxa with a minimum prevalence of 10% and a minimum abundance of 0.01% were considered. P-values were adjusted for multiple comparisons using the Benjamini-Hochberg (FDR) method. The effect size (coefficient) reflects the log2-fold change in abundance relative to the 2 h reference.

**Figure 4.**
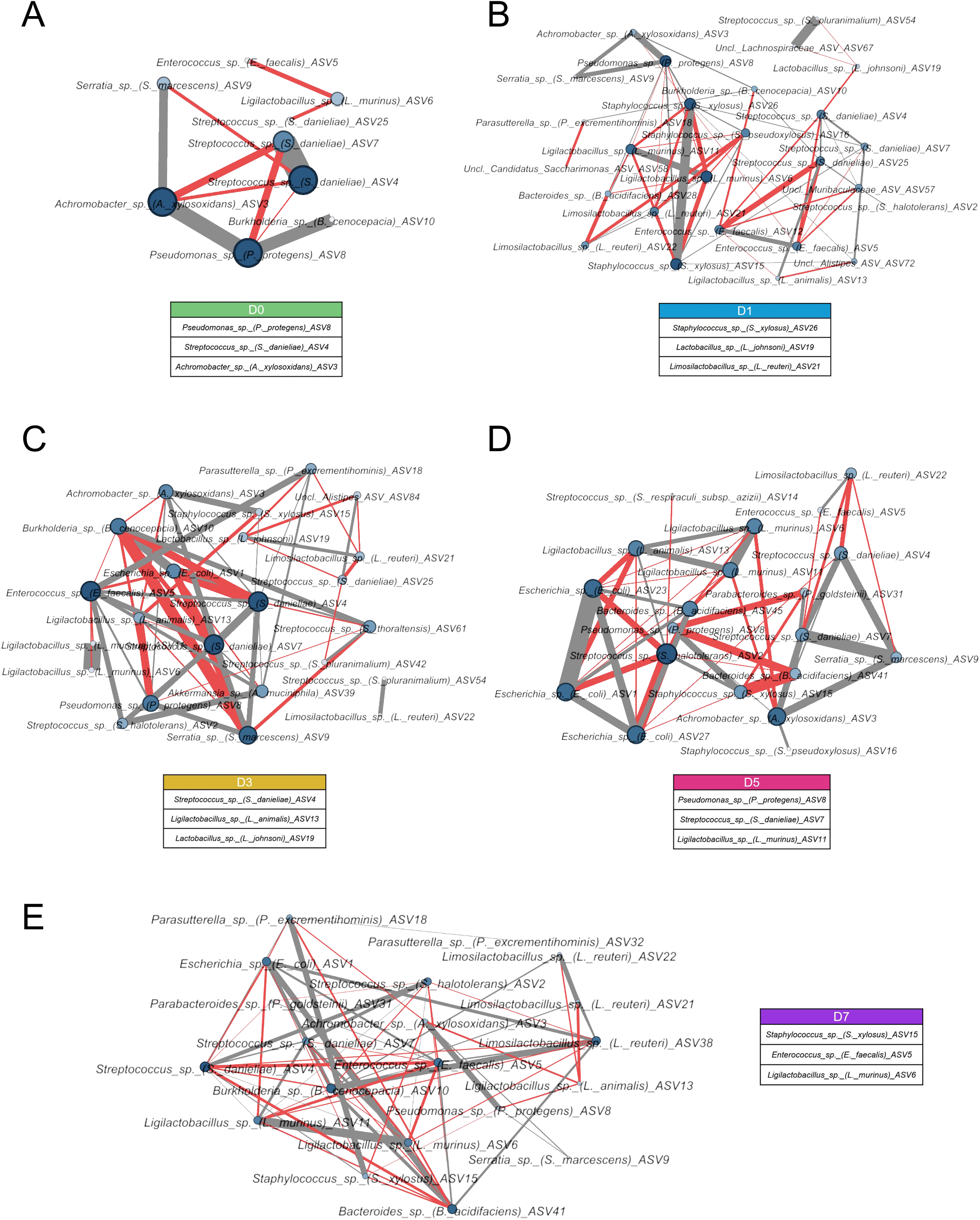
Bacterial co-occurrence networks across the timeline of ligature-induced periodontitis. Networks are shown for Day 0 (A), Day 1 (B), Day 3 (C), Day 5 (D) and Day 7 (E) post-ligature placement. The thickness of the edges indicates the strength of the Spearman correlation (|r| > 0.6), with grey representing positive correlations and red representing negative correlations. The size of each node (ASV) and the color darkness represent the proportionality to the number of connections (degree). For each network, the top 3 ASV with the highest number of connections are labeled. For all networks, only taxa present in at least 30% of the samples were included.

At D1, a compositional shift was observed, marked by the emergence of new taxa and the establishment of a distinct microbial profile. These included *Streptococcus_sp._(S._halotolerans)_ASV2, Enterococcus_sp._(E._faecalis)_ASV5, and Ligilactobacillus_sp._(L._murinus)_ASV6,* the latter being present at lower relative abundance. This stage was also characterized by a reduction in HAT (Fig. 3A). However, HAT remained enriched compared to later stages, such as D3 (Fig. S3B).

By D3, a dysbiotic state was established, characterized by two distinct microbial profiles: one primarily driven by *S._halotolerans_ASV2* and another by *Escherichia_sp._(E._coli)_ASV1* (Fig. 3A). This time-point also featured the presence of *S._danieliae_ASV4/7, L._murinus_ASV6/11, and E._faecalis_ASV5*, albeit at lower relative abundance (Fig. 3A). From D3, few discriminant features were detected between groups, with no significant differences observed between D5 and D7 (Fig. S3C). Overall, these time points were characterized by a decline in HAT, alongside the emergence and dominance of taxa such as *E._coli_ASV1, S._halotolerans_ASV2, L._murinus_ASV6/11, and E._faecalis_ASV5*, collectively defining a group of periodontitis-associated taxa (PAT) (Fig. 3A, S3C, S4). Notably, a reciprocal and mutually exclusive pattern was observed between *S._halotolerans_ASV2* and *E._coli_ASV1*, in which the dominance of one coincided with the depletion of the other (Fig. 3A). These findings reveal an early decrease of HAT that precedes the establishment of a dysbiotic state marked by different dominance patterns between PAT.

Given the dominant bacterial species on each day, probes were designed for Combinatorial Labeling and Spectral Imaging Fluorescence in situ Hybridization (CLASI-FISH) to visualize them in the ligature. On D0, very few bacterial species were detected, consistent with the lowest bacterial load values. However, HAT such as *A. xylosoxydans* were observed (Fig. 2B, upper middle; S5). On D1, bacteria such as *S. danielae* and *S. halotolerans* were detected in the ligature, either together or in separate clusters (Fig. 2B, upper right). On D3, the proportion of *E. coli* increases as biofilm accumulates around the ligature (Fig. 2B), with *S. halotolerans* (Fig. S5). Subsequently, from D3 onward, the ligature biofilm can be dominated by either *S. halotolerans* or *E. coli*, but not by both simultaneously, thereby establishing two distinct, spatially segregated microbial profiles, indicating mutually exclusive abundance patterns. (Fig. 2B, lower left; S5). Importantly, despite the fact that *S. danieliae* and *A. xylosoxydans* are not dominant taxa in the established dysbiosis profiles (D7), they are still present in lower proportions. Taken together, these observations highlight the development of structurally organized and spatially distinct communities during disease progression. These spatial patterns were consistent with the relative abundance data (Fig. 3A). Additionally, CLASI-FISH revealed an increased bacterial accumulation over time, which was complemented by scanning electron microscopy (Fig. S6).

Additionally, temporal trend analysis of the ten most dynamic taxa revealed a clear divergence in successional patterns (Fig. 3C). The relative abundances of certain PAT, such as *E._coli_ASV1* and *S._halotolerans_ASV2,* increased progressively over time starting D1 onward. Conversely, HAT, such as *A._xylosoxidans_ASV3* and *S._danieliae_ASV4/7*, exhibited a consistent decline throughout the experimental period.

### Network analyses reveal increased complexity featuring negative correlations during dysbiosis establishment

Network restructuring reveals increased complexity and likely antagonistic interactions during dysbiosis establishment (Fig. 4). To investigate the temporal dynamics of microbial interactions, a co-occurrence network was constructed for each day, from baseline to D7. For each network, the three taxa (ASVs) with the highest number of connections, regardless of their nature (positive or negative), were identified as key influencers (Fig.4). An increase in network complexity was observed from D0 (Fig. 4A) to D1 (Fig. 4B). The D0 network comprised only 8 nodes (taxa) and 11 edges (correlations). From D1 onward, all networks were composed of 20 or more nodes, with D1 exhibiting the highest number (26 nodes), followed by D3 (23) (Fig. 4C), and D5 (Fig. 4D) and D7 (20 each) (Fig. 4E).

In the D0 network, the three most connected and abundant taxa were *P._protegens_ASV8, S._danieliae_ASV4*, and *A._xylosoxidans_ASV3*. This early community was characterized by negative correlations, including interactions among the aforementioned species. As dysbiosis progressed, *P._protegens_ASV8* and *A._xylosoxidans_ASV3* maintained their influencer status across all days, with a predominance of positive correlations despite decreases in their relative abundance over time. Notably, *S._danieliae_ASV4* exhibited a consistent tendency for antagonistic (negative) correlations throughout the time series, particularly with *E._coli_ASV1* and *E._faecalis_ASV5*. In contrast, *S._danieliae_ASV7* has more positive correlations. A similar negative correlation to ASV4 was observed for *S._danieliae_ASV25*, which disappeared from the network after D3.

The microbial taxa that emerged after D0 exhibited distinct and consistent correlation patterns. *E._faecalis_ASV5* and *L._murinus_ASV6/11*, both of which increased in relative abundance from D1 onward, were constant network members and maintained a stable pattern of negative correlations with each other. *E._coli_ASV1*, associated with a dysbiotic profile in the relative abundance plots, first appeared as a network component at D3. *S._halotolerans_ASV2*, central to the other dysbiotic profile, emerged at D1 and significantly increased its number of positive correlations from D3 onward. Other persistent members included *S._marcescens_ASV9* with primarily positive correlations, *Staphylococcus_sp._(S._xylosus)_ASV15* (primarily positive correlations), and *Ligilactobacillus_sp._(L._animalis)_ASV13* (primarily negative correlations); the latter two taxa appeared at D1 and persisted through all subsequent time points.

### The established dysbiotic microbiome exhibits a predicted functional profile compatible with increased bacterial virulence

Functional prediction analysis revealed metabolic restructuring of the ligature microbiome throughout dysbiosis progression. Pairwise comparisons of D0 against each subsequent day (D1, D3, D5, D7) highlighted significant differential abundance in key KEGG pathways (ALDEx2, BH-corrected Wilcoxon rank test) (Fig. 5A-D). From the initial comparison (D0 vs. D1), there is a significant change in the microbial functional predicted profile (Fig. 5A). Although the relative abundances of the top-enriched functions appeared similar, there were pathways differentially represented. Pathways enriched in D0 were dominated by those supporting bacterial motility (bacterial chemotaxis, flagellar assembly), virulence (bacterial secretion systems), and metabolism. In contrast, D1 showed a higher relative abundance of pathways involved in secondary metabolite biosynthesis. The difference in predicted function increased by D3 (D0 vs. D3) (Fig. 5B). Functions related to cellular motility remained significantly elevated in D0. However, D3 exhibited a significant increase in protein export, a pathway critical for virulence factor secretion. Notably, various core metabolic pathways (carbohydrate metabolism, metabolism of cofactors and vitamins) were predominantly more abundant in the D0 community. A shift in this trend was observed by D5 (D0 vs. D5) (Fig. 5C). While bacterial chemotaxis was enriched in D5, protein export decreased relative to earlier points. Concurrently, pathways for xenobiotic biodegradation and metabolism became significantly elevated in D5. By D7 (D0 vs. D7), the predicted functional profile of the established dysbiotic microbiome consolidated into a distinct state consistent with enhanced pathogenic potential (Fig. 5D). Bacterial chemotaxis was significantly higher in D0, whereas protein export was again significantly increased in D7. Furthermore, D7 showed enrichment across diverse metabolic pathways, most notably galactose metabolism, lipid metabolism, cofactor and vitamin metabolism, and xenobiotic biodegradation and metabolism.

**Figure 5.**
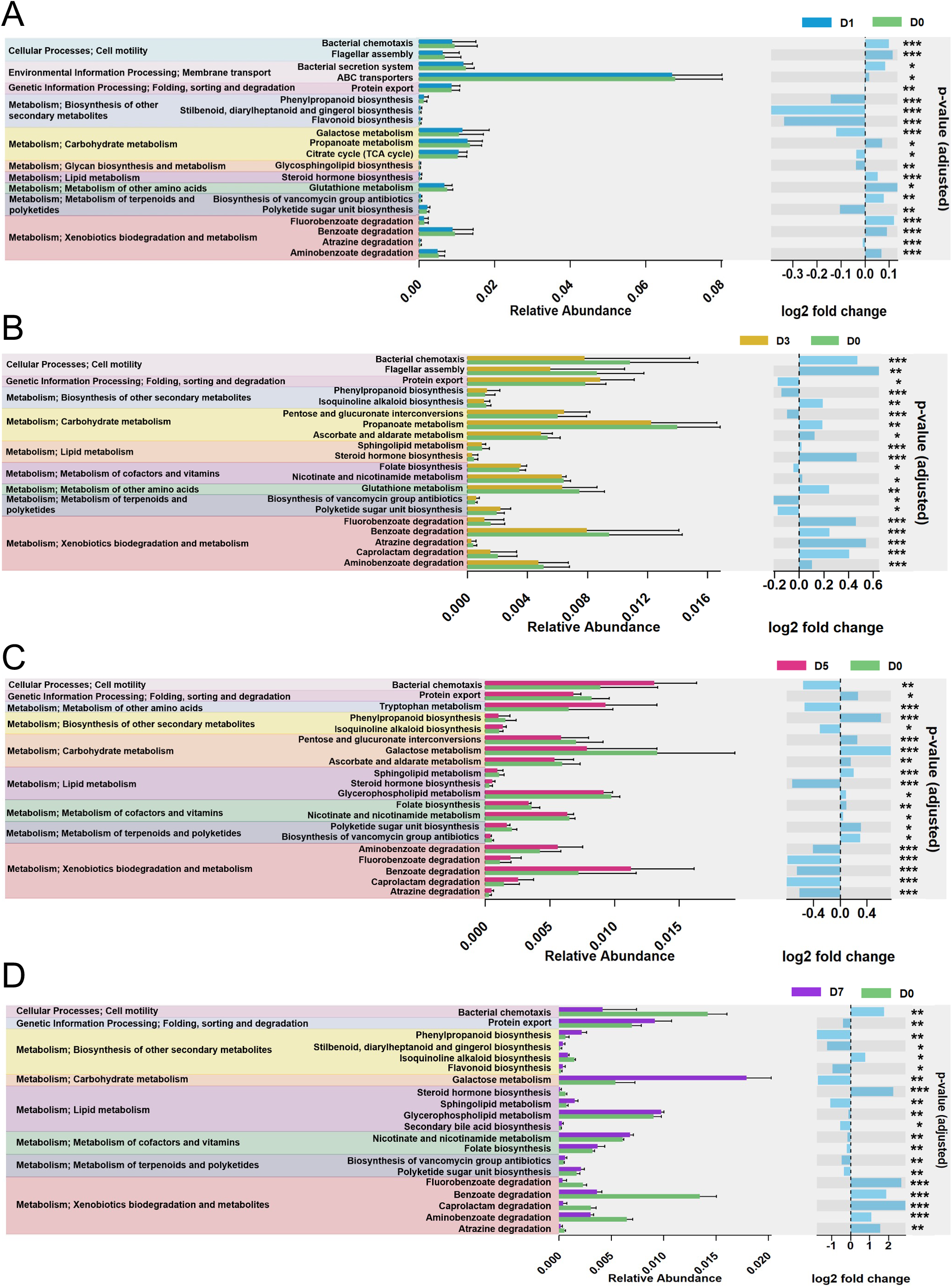
Differential functional profile predictions of the ligature-induced microbiome across time points. Functional predictions were generated from 16S rRNA data using PICRUSt2 and analyzed for differential abundance. Pathways were annotated using the KEGG database. (A-D). Bar plots representing the top 20 most abundant KEGG pathways with significant differences for each comparison: Day 0 vs. Day 1 (A), Day 0 vs. Day 3 (B), Day 0 vs. Day 5 (C), and Day 0 vs. Day 7 (D). The fold-change plot adjacent to each bar plot indicates the direction and magnitude of the differential change between the two compared time points. Differential abundance analysis was performed using the ALDEx2 method (Wilcoxon rank test) with Benjamini-Hochberg (BH) correction for multiple comparisons. Significance levels are denoted as follows: *p < 0.05, **p < 0.01, ***p < 0.001.

### Variable association modeling integrates microbial and host features of disease progression, revealing the importance of health-associated taxa

Finally, we identified significant associations between microbial features and host phenotypes using an analysis tool specifically designed to detect these comparisons (MaAsLin 3). Several ASVs, including *A. xylosoxidans*, *S. danieliae*, *P. protegens*, *S. marcescens*, and *B. cenocepacia*, showed negative abundance associations with total bacterial load (Fig. 6A; Fig. S7). The same taxa exhibited consistent negative associations with the percentage of immune (CD45^+^) cells: *A. xylosoxidans*, *S. danieliae* ASV4 and ASV7, *P. protegens*, *S. marcescens*, and *B. cenocepacia* (Fig. 6B; Fig. S8). In contrast, *E. faecalis* showed a positive abundance association with CD45^+^ cells, progressively increasing until reaching a plateau (Fig. 6B; Fig. S8). Regarding collagen content, higher collagen percentages were positively associated with *A. xylosoxidans*, *P. protegens*, *S. marcescens*, and *B. cenocepacia* (Fig. 6C; Fig. S9). All associations described above were driven by abundance rather than prevalence; no significant prevalence associations were detected for any of the host features examined. Furthermore, no significant associations, either in abundance or prevalence, were observed for alveolar bone loss or TRAP^+^ cells (Fig. S10). Collectively, these findings reveal that health-associated taxa, including *A. xylosoxidans, S. danieliae, P. protegens, S. marcescens*, and *B. cenocepaci*a, were negatively associated with disease-associated features (total bacterial load and CD45⁺ cells) and positively associated with collagen content, consistent with their lower relative abundance levels during experimental periodontitis.

**Figure 6.**
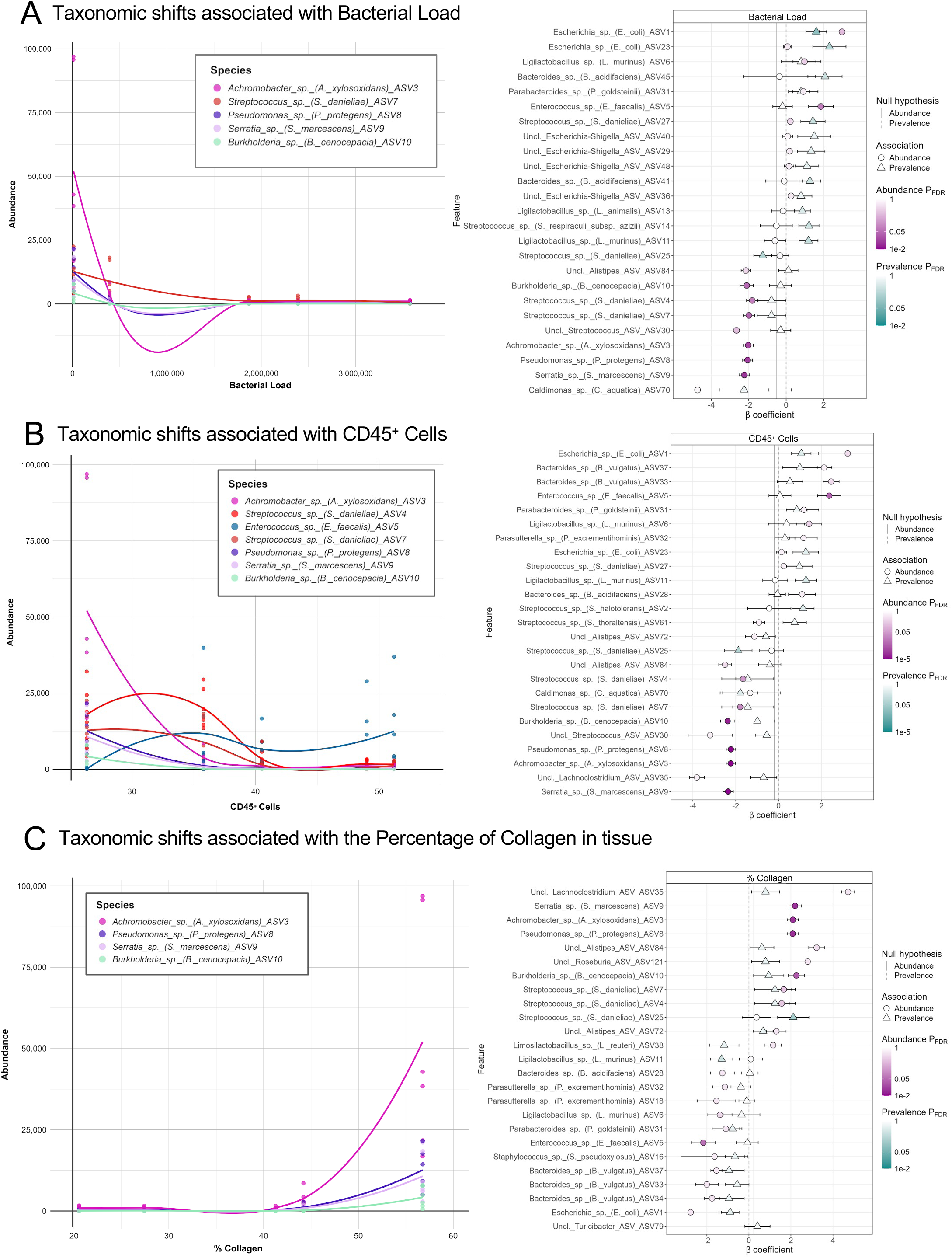
Significant microbial associations with alveolar bone loss, bacterial load, CD45^+^ cells, and collagen. (A) Taxonomic shift associated with bacterial load and forest plot of top microbial features associated with bacterial load. (B) Taxonomic shift associated with CD45^+^ Cells and forest plot of top microbial features associated with CD45^+^ Cells. (C) Taxonomic shift associated with percentage of collagen and forest plot of top microbial features associated with percentage of collagen. Trends of significant taxa, identified by a MaAsLin 3. The analysis was run using a Linear Model (LM) on TSS-normalized and LOG-transformed data. Only taxa with a minimum prevalence of 10% and a minimum abundance of 0.01% were considered. P-values were adjusted for multiple comparisons using the Benjamini-Hochberg (FDR) method. Forest plot points represent effect sizes (coefficients) from MaAsLin 3 models, with horizontal lines indicating 95% confidence intervals. Features are ordered by significance (top = most significant). Colors distinguish abundance (blue) versus prevalence (red) associations. Negative coefficients indicate ASVs that decreased with the corresponding variable; positive coefficients indicate ASVs that increased with the corresponding variable. Labels depict the name of each ASV.

Our findings reveal a temporal sequence of microbial and tissue events in which the decrease in HAT precedes and correlates with inflammatory responses and tissue destruction. At D1, a shift in microbiome structure occurs, followed by the establishment of a stable dysbiotic profile by D3. CD45⁺ cell infiltration increases at D1, preceding the rise in TRAP⁺ cells at D3 and tissue destruction by D5.

## Discussion

In this study, histological and microbiological events were analyzed at longitudinal experimental timepoints to provide an integrated view of experimental periodontitis establishment in the LIP model, incorporating both compositional and structural features of the microbiome. Under steady conditions, low bacterial load and a microbiome enriched in HAT were observed, with no evidence of inflammation or tissue destruction, resembling periodontal health ^24,25^. By D1, the microbiome already shows early compositional and structural changes, indicating a rapid transition from homeostasis. These changes are sufficient to trigger an initial immune response, as shown by the increase in immune cells. This increase may reflect early neutrophil recruitment, as a significant rise in these cells has been reported in periodontal tissues at comparable time points^17,18^. Notably, this phase has also been associated with increased expression of pro-inflammatory cytokines and activation of neutrophil-related inflammatory pathways ^16,26^, suggesting that the establishment of an initial inflammatory response may be driven by neutrophil recruitment.

At D3, the microbiome reaches a dysbiotic state, characterized by a higher bacterial load and a significant increase in TRAP⁺ cells, consistent with previous reports ^18,22^. This increase indicates the accumulation of cells with osteoclastogenic potential, which will then drive bone resorption. In fact, this stage may represent a critical window for tissue destruction, as neutrophil depletion at D3, but not at later stages, has been shown to prevent immune-driven bone loss ^27^. By D5 and D7, the microbial dysbiotic state appears established, with elevated levels of CD45⁺ and TRAP⁺ cells relative to baseline, also including significant collagen and alveolar bone loss. This observation is consistent with previous reports indicating that bone loss becomes detectable around days 4 and 5 post-ligation, accompanied by collagen degradation ^14,17,23^. These histological changes appear to be locally driven by the bacterial stimulus at the ligature, as no bacterial invasion into periodontal tissues was observed, suggesting that disease progression in this model is primarily mediated by host responses to the biofilm rather than by direct bacterial infiltration ^28–30^. However, bacterial presence in deeper tissues cannot be completely excluded, as ultrastructural analyses have revealed bacteria within osteocyte lacunar spaces during the LIP model ^18^.

In agreement with Marchesan et al. (2023) and Ribeiro et al. (2022), a progressive decrease in alpha diversity and a clear ecological succession were observed. Importantly, the most pronounced microbiome shift occurs at D1, resembling the transition observed in human gingivitis, in which health and dysbiosis-associated species coexist ^5,31^. In contrast to gingivitis, the murine microbiome does not exhibit increased species richness; instead, it is characterized by shifts in the relative abundance of pre-existing species ^5^. Beta diversity analyses also identified D1 as a transitional state preceding the establishment of dysbiosis, which becomes stable from D3 onward, as indicated by the substantial overlap among D3, D5, and D7 clusters. Differential bacterial abundance analyses confirmed this, revealing fewer discriminative taxa between D3 and D5 and none between D5 and D7. Together, these findings highlight how the early decrease in HAT and the expansion of PAT drive the microbiome toward a stable dysbiotic state that may contribute to disease persistence.

Our data identified *S. danieliae* as a taxon associated with periodontal homeostasis in mice, a finding consistent with the bioinformatic reanalysis by Arce et al. (2022). However, our study reveals a consortium of commensal taxa that exhibits a previously undescribed temporal dynamic in relation to the host response. This group, comprised of ASVs that are identified as *A. xylosoxidans*, *P. protegens*, *S. marcescens*, and *B. cenocepacia*, was first identified through correlation network analysis and subsequently validated as taxa of interest. Similarly to what occurs in humans, these HAT persist during the transition toward dysbiosis, albeit experiencing a dramatic decrease in relative abundance. This stability aligns with our alpha-diversity analyses, suggesting a reconfiguration of community structure rather than the loss of HAT. This ecological succession, characterized by an expansion of pathobionts, enhances community complexity as disease progresses. Consistent with observations in human periodontitis and peri-implantitis, network complexity increases as dysbiosis becomes established, reflecting the reorganization of microbial interactions during disease progression ^5,32^.

The ecological fitness of HAT is likely to stem from a suite of highly conserved competitive and metabolic mechanisms. Certain strains of *A. xylosoxidans* have high biofilm-forming capacity ^33^ and outcompete other bacteria by inhibiting initial adhesion, extracellular matrix production, disrupting secretion systems, and iron uptake ^34^. Similarly, *P. protegens* and *B. cenocepacia* utilize iron sequestration to compete within the niche, limiting resource availability for other microorganisms without compromising host integrity ^35–37^. This is further complemented by the production of antimicrobial, antifungal, and immunomodulatory compounds described in *P. protegens*, *S. marcescens*, and *B. cenocepacia* ^38–41^. These HAT likely possess intrinsic competitive traits that enable them to stabilize the ecological niche under homeostatic conditions. However, upon ligature-induced dysbiosis, the relative abundance of these HAT declines, creating an opportunity for pathobionts such as *E. faecalis* and *E. coli* to expand.

Additionally, a critical divergence in healthy microbiome composition was observed: while human commensals are predominantly Gram-positive ^5^ our data confirm that HAT in mice are mostly Gram-negative, aerobic, and motile ^42,43^. These traits likely explain the functional signature detected via PICRUSt2, which shows an enrichment of motility-related pathways at baseline (D0).

Mice are naturally resistant to periodontitis ^44^, yet this resistance is likely not due to the absence of periodontopathic bacteria, since their indigenous microbiota can trigger disease ^45^. Rather, we propose that the Gram-negative consortium actively constrains pathobiont expansion under homeostatic conditions. The ligature disrupts this suppression by increasing bacterial biomass and reducing oxygen tension, thereby favoring facultative anaerobes such as *E. coli* and *E. faecalis*.

Among HAT, the case of *S. danieliae* is particularly compelling. Despite being Gram-positive and not following the dynamics of the Gram-negative consortium, its phylogenetic proximity to *S. sanguinis*, which constitutes a health-associated species in the human subgingival microbiome, suggests a significant ecological role. This similarity transcends taxonomy: in both human gingivitis and the early transition stage (D1) in mice, this taxon maintained stable abundances, emerging as a differential species only when contrasting health against established periodontitis ^5^. Furthermore, in humans, *S. sanguinis* is essential for microbiome assembly, acting as a primary colonizer and playing a key role in biofilm formation ^46^, while also facilitating the adhesion and colonization of pathobionts such as *P. gingivalis* ^47^. A similar phenomenon has been reported for *S. danieliae*, which possesses the ability to co-aggregate with *F. nucleatum* and specific strains of *P. gingivalis* ^43^, suggesting that *S. danieliae* may be considered as the murine functional equivalent of *S. sanguinis*. This implication further underscores the LIP model’s relevance for understanding the microbial transition from periodontal health to disease. Although taxonomical identity between humans and mice varies, their roles in the overall microbial community might be conserved.

The decline of HAT, coupled with the expansion of *E. coli, E. faecalis, S. halotolerans* and *L. murinus*, reflects a broader shift toward a dysbiotic state that becomes functionally distinct by D7. *E. faecalis* emerges as one of the earliest taxa to increase in abundance, serving as a key discriminant marker between baseline (D0) and all subsequent time points (D1, D3, and D7), consistent with previous reports ^15^. Furthermore, the expansion of *E. faecalis* positively correlates with increased numbers of immune cells. This late-stage microbiome is characterized by the enrichment of pathways related to protein export and xenobiotic metabolism, aligning with the upregulation of TLR-signaling pathways reported by Bao et al. (2019) and the stabilization of inflammatory infiltrates observed by Tsukasaki et al. (2018). Collectively, these findings position *E. faecalis* as a discriminative microbial marker distinguishing periodontal health from established disease in the LIP model. The visualization of ligature-associated communities using CLASI-FISH, suggests that the ligature itself appears to serve as the physical support for biofilm formation in the LIP model, potentially bypassing the need for filamentous bacterial scaffolding. In humans, the spatial architecture of supragingival plaque is structured by filamentous bacteria such as *Corynebacterium* and *Fusobacterium*, which form a scaffold that organizes other taxa into radially arranged consortia ^48^. In contrast, the murine microbiome lacks such filamentous organizers. Instead, bacteria seem to organize into discrete, spatially segregated clusters, with *E. coli* and *S. halotolerans* never observed in close proximity. This mutually exclusive colonization pattern aligns with our abundance data, which show that these two species do not co-dominate. Spatial imaging revealed a distinctive biofilm architecture in the murine LIP model, shaped by the ligature as an exogenous scaffold and characterized by ecological exclusion between key dysbiotic taxa.

Finally, the mutual exclusivity observed between *E. coli* and *S. halotolerans* warrants particular attention. Although *S. halotolerans* remains poorly characterized, it is defined as a Gram-positive coccus capable of acid production through sugar fermentation ^49^. Given that many *Streptococcus* spp. produce hydrogen peroxide (H_2_O_2_) as a competitive mechanism ^50–53^, we hypothesize that *S. halotolerans* may employ a similar strategy. The synergistic effect of lactic acid and H_2_O_2_ has been shown to inactivate *E. coli* by generating hydroxyl radicals that damage DNA and cellular membranes ^54,55^. We can hypothesize that this antagonistic interaction likely explains why these two species cannot simultaneously dominate the same dysbiotic profile, further highlighting the role of niche competition in shaping subgingival community types.

While diverse dysbiotic profiles in mice have been noted previously ^15^, the depth of this heterogeneity remains underexplored. Distinct microbial signatures have been identified depending on laboratory setting, displaying inter-individual variation, yet bone loss remained a consistent outcome, a pattern of inter-individual variation also reported in human periodontitis ^5^. We argue that extrinsic environmental factors in murine models (handler interactions, water pH, husbandry conditions) may emulate the inherent complexity of the human microbiome ^56,57^. A reflection of this heterogeneity is that only four consistent markers of murine periodontitis have been identified ^15^, of which *E. faecalis* was the only shared marker in our study. Importantly, methodological differences, such as defining OTUs versus higher-resolution ASVs may account for these discrepancies, as previously unclassified taxa likely correspond to the specific members identified here.

Beyond characterizing microbial succession, our study differs from previous contributions in the field by directly associating microbiome composition with host tissue parameters. We observed that HAT abundance was positively associated with collagen percentage and negatively associated with both total bacterial load and immune cell infiltration. Consistent with the competitive and immunomodulatory traits of HAT described above, the negative HAT-CD45⁺ and positive HAT–collagen associations suggest a protective role. As previously mentioned, during the early stages of disease, the loss of HAT dominance within the microbial community is associated with increased immune cell infiltration, including neutrophils and other immune populations that can release matrix metalloproteinases and collagenases, thereby contributing to collagen degradation ^58,59^. In contrast, the pathobiont *E. faecalis* displayed the inverse relationship, correlating positively with CD45⁺. This association is mechanistically plausible, as *E. faecalis* is known to activate pro-inflammatory cytokine production and can survive intracellularly within macrophages, thereby sustaining immune cell recruitment ^60^. This finding positions *E. faecalis* as a driver and marker of dysbiosis. While our correlational approach does not allow us to infer causality or determine whether microbial changes drive host responses or vice versa, it established robust links between specific taxa and quantifiable host parameters. Importantly, they suggest that HAT may contribute to tissue homeostasis not only by constraining pathobiont expansion but also by correlating with preserved collagen architecture and reduced immune infiltration.

This multifaceted framework was achieved by integrating histological analyses with advanced microbiome bioinformatics and spatial imaging techniques, thereby highlighting the added value of our integrative approach. This strategy allowed us to detect that the D1 microbiome mirrors human gingivitis, whereas by D3 the community has already reached a dysbiotic state. Moreover, CLASI-FISH imaging revealed that bacteria organized into discrete, spatially segregated clusters, with *E. coli* and *S. halotolerans* never observed in close proximity, featuring a mutually exclusive abundance pattern inaccessible to sequencing alone. This spatial resolution also revealed that HAT forms a spatially organized, Gram-negative consortium that could maintain ecological stability under homeostatic conditions. Our results, corroborated by sequencing and histological data, show that the ecological shift to dysbiosis (D1-D3) is the decisive event driving the subsequent destructive cascade. This integrated framework not only validates the ligature-induced periodontitis model but also identifies the specific cellular and microbial windows where therapeutic intervention might be most effective. Ultimately, it suggests that future intervention strategies should focus on preserving or restoring the eubiotic state, rather than targeting the variable members of the dysbiotic community.

## Materials and Methods

### Mouse strains and experimental periodontitis

C57BL/6N mice (10–14 weeks old, with equal numbers of males and females) were bred and maintained under specific pathogen-free (SPF) conditions at the Faculty of Dentistry, University of Chile. All procedures were approved by the Institutional Animal Care and Use Committee (IACUC) and comply with ARRIVE guidelines. LIP was performed as previously described (Abe & Hajishengallis, 2013) by placing a 5-0 silk ligature around the upper left second molar, with the contralateral molar serving as an internal control. Mice were euthanized at D0 (2 h), D1 (24 h), D3 (72 h), D5 (120 h), or D7 (168 h) post-ligation. Periodontal tissues (gingiva, bone, and teeth) were collected, fixed in 4% paraformaldehyde, decalcified in 5% EDTA, and embedded in paraffin for histological analysis. In parallel, an additional cohort was used for morphometric assessment of alveolar bone loss, and ligatures were collected in TE buffer for further microbial sequencing analyses.

### Alveolar Bone Loss Measurements

Skulls were boiled to facilitate soft-tissue removal and were subsequently defleshed. Maxillae were then treated with 3% hydrogen peroxide, stained with 1% methylene blue, and photographed using a Zeiss Stemi 508 stereomicroscope with an Axiocam ERc 5s camera (3.5X). Alveolar bone height was measured in ImageJ by assessing the distance from the cementoenamel junction (CEJ) to the alveolar bone crest (ABC) at six predefined sites per sample. Bone loss was calculated as the difference in CEJ–ABC distance between ligated and contralateral unligated sides ^6,14^.

### Histological processing and staining

Tissue sections (3–5µm) were deparaffinized in xylene and rehydrated through graded ethanol solutions (100%-95%-70%) to distilled water. Hematoxylin and eosin (H&E) staining was performed using Gill’s hematoxylin (10min) followed by eosin (3 min). For immunohistochemistry (IHQ), antigen retrieval was carried out using citrate buffer (pH 6), endogenous peroxidase activity was blocked with 3% H₂O₂ in methanol, and sections were incubated with casein prior to overnight incubation at 4°C with anti-CD45 primary antibody (1:200; catalog 14045182, eBioscience) followed by an incubation with anti-rat secondary antibody (1:200; catalog 712065153, Jackson Immuno Research) at room temperature for 1 hour. Detection was performed using the VECTASTAIN Elite ABC HRP system and DAB substrate (ImmPACT DAB Substrate, Vector Laboratories), followed by hematoxylin counterstaining. CD45⁺ cells were quantified using FIJI (ImageJ) and expressed as the percentage of positive cells relative to the total number of cells within the lamina propria of the interproximal area between the first and second molars. Tartrate-resistant acid phosphatase (TRAP) staining was performed using a commercial kit (Leukocyte Acid Phosphatase (TRAP) Kit MERCK) according to the manufacturer’s instructions. TRAP-positive cells were identified by purple/brown cytoplasmic staining and were quantified by counting positively stained multinucleated cells within the periodontal area of interest under light microscopy. Masson’s trichrome staining was used to assess collagen. Sections were sequentially stained with Weigert’s iron hematoxylin and Biebrich scarlet–acid fuchsin, followed by incubation in phosphomolybdic–phosphotungstic acid, subsequent staining with aniline blue, and a final step in 1% acetic acid. Collagen quantification was performed with FIJI by selecting the connective tissue area, excluding bone and dental structures, followed by color deconvolution (H PAS) and thresholding to isolate collagen fibers. The analysis was expressed as the percentage of collagen relative to the total analyzed area. Brown and Brenn staining was performed to visualize bacteria in the ligature and surrounding tissues. Sections were sequentially stained with crystal violet, treated with iodine solution, briefly decolorized with acetone, counterstained with fuchsin, and exposed to a picric acid–acetone solution prior to clearing. Following all staining procedures, sections were dehydrated through graded alcohols, cleared in xylene, and mounted with Entellan (MERCK). All histological sections were examined under an optical microscope (Axiolab 5, Zeiss), and representative images were captured at 20X magnification using a digital camera (Axiocam 208 color, Zeiss).

### DNA Isolation and total bacterial load assessment

The DNA was extracted from the ligature samples using a modified protocol of the DNeasy Blood and Tissue kit (Qiagen), which involved lysozyme and proteinase K treatment as previously described ^61^.

The total bacterial load in the ligature was measured by qPCR using 16S rRNA universal primers ^24,62^. Genomic DNA from *Fusobacterium nucleatum* ATCC 10953 was extracted and used to construct standard curves with a known number of 16S rRNA gene copies (ranging from 10^2^ to 10^8^). These standard curves were used to determine the number of 16S rRNA gene copies in each sample.

### 16S rRNA Sequencing and Bioinformatic Processing

DNA from ligature samples was sequenced for the V4 region of the 16SrRNA gene using Illumina MiSeq. Amplicon sequence variants (ASVs) were inferred using DADA2 version 1.16 ^63^ in the R environment. Briefly, this involved quality filtering and trimming, error rate estimation, dereplication, sample inference, and merging paired-end reads. Chimeric sequences were identified and removed, resulting in an ASV table. Taxonomy was assigned to the final ASVs using the SILVA reference database (v138.2) ^64^.

### Microbiome Data Analysis and Statistics

All subsequent microbiome data analysis and statistical computations were performed in R. The resulting ASV table, taxonomy table, and sample metadata were imported and assembled into a phyloseq object (version 1.48.0, ^65^). Relative abundance profiles were calculated. Alpha diversity indices (which measure ecological variation within microbial communities), were estimated including observed ASVs, Shannon, and Inverse Simpson indices. Beta diversity, which compares microbial community composition between samples, was assessed using multiple distance metrics: Bray-Curtis, Jaccard, Yue & Clayton (ThetaYC), and phylogenetic-based Unweighted and Weighted UniFrac distances. Principal coordinate analysis (PCoA) was performed for ordination, and permutational analysis of variance (PERMANOVA) with 9,999 permutations was used to test for significant differences between groups. Differential abundance analysis was performed using Linear discriminant analysis Effect Size (LEfSe) ^66^. Statistical significance was assessed using the Kruskal-Wallis test (p <0.05), followed by pairwise Wilcoxon rank-sum test (p <0.05). Features with a logarithmic LDA score >2.0 were considered differentially represented. Co-occurrence analyses were conducted based on Spearman correlation values of ASV-level taxa with the package Netcomi ^67^, and the predictive functional potential of the microbial communities was explored using PICRUSt2 ^68^.

To identify taxa associated with host phenotypes, we performed longitudinal association analyses using MaAsLin 3 ^69^ with TSS-normalized and log-transformed abundance data, setting D0 as the reference. Taxa with ≥10% prevalence and ≥0.01% abundance were included. Effect sizes represent log2-fold changes relative to D0, and p-values were adjusted using the Benjamini-Hochberg FDR method. Mice were grouped by time point, and the mean phenotype value for each group was assigned to all mice within that group. Associations with q <0.05 were considered significant.

### CLASI-FISH

Species-specific probes for *Streptococcus thoraltensis* (now *Streptococcus halotolerans*)*, Streptococcus danieliae, Escherichia coli, Achromobacter xylosoxidans*, and *Ligilactobacillus murinus* were designed in silico using the DECIPHER software ^70^. Probes targeting the 23S rRNA were designed for *S. danieliae* and *S. thoraltensis*, whereas probes targeting the 16S rRNA were designed for the other species. All oligonucleotides were designed to be 18-22 nucleotides in length, demonstrate 100% specificity to their intended target with zero off-target hits, and possess an estimated overall change in Gibb’s free energy upon binding between -12 and -14 kcal/mol. A two-step fluorescence in situ hybridization (FISH) assay was performed. Briefly, fixed cells were first hybridized with an unlabeled encoding probe containing a species-specific targeting sequence and a readout sequence. After stringent washes, a fluorophore-labeled readout probe was hybridized to readout sequence ^71^. Spectral imaging was performed on a Zeiss LSM 980 confocal microscope with a 63x/1.4 NA objective. Linear unmixing was performed using spectra extracted from reference images. Final image processing, including Z-projection, segmentation to remove tissue autofluorescence, and channel merging, was conducted in FIJI to generate the final composite images ^72^.

### Scanning Electron Microscopy (SEM)

The tissues with intact biofilm were fixed in 2.5% glutaraldehyde in 0.1 M sodium cacodylate buffer (pH 7.4) for 24 h at 4°C, washed in cacodylate buffer, and dehydrated through a graded ethanol series. Samples were critical-point dried using liquid CO₂, mounted on aluminum stubs, sputter-coated with gold-palladium, and imaged with a scanning electron microscope (Jeol JSM-IT300LV, JEOL USA Inc) operated at 20.0 kV under high vacuum.

## Data availability

The raw sequencing data generated and analyzed in this study have been deposited in the NCBI BioProject database under accession number PRJNA1457214.

## Acknowledgments

The authors thank Matías Morales and Rocio Orellana (Project Fondequip EQM130076), for their technical support. This work was funded by ANID – FONDECYT Regular (1231350 to ND and 1231728 to LA) and by NIH/NIDCR (R01DE031213 to AMV and LA). Additionally, it was supported by the doctoral scholarship from the University of Chile (to N.E) and by scholarships from ANID, provided by the Chilean government (2024-21240601 to J.E.-A, 2025-21250805 to M.I.M.-S, 2022-21221003 to M.A and 2023-21231392 to D.S). We would like to specially thank the Experimental Platform from the Faculty of Dentistry, University of Chile

## Conflict of interests

The authors declare no conflict of interest with respect to the research, authorship, and/or publication of this article.

## Contributions

N. Endo and J. Espinoza-Arrue contributed to data acquisition, analysis, interpretation and drafted the manuscript. M. Arce, N. Traver, MI. Muñoz-Sepúlveda, D. Sansores-España, V. Olmedo, C. Moreno, J. Canelo contributed to data acquisition, analysis, and critically revised the manuscript. AM. Valm and M. Reyes supervised data acquisition, contributed to data interpretation, and critically revised the manuscript. L. Abusleme and N. Dutzan contributed to conception and design, data acquisition, analysis, interpretation, and drafted the manuscript. All authors gave final approval and agreed to be accountable for all aspects of the work.

## Notes

### Competing Interest Statement

The authors have declared no competing interest.

