## Supplementary figures for "“A decrease in specific health-associated commensals is linked to progressive periodontal tissue destruction independent of dysbiotic community profiles”"

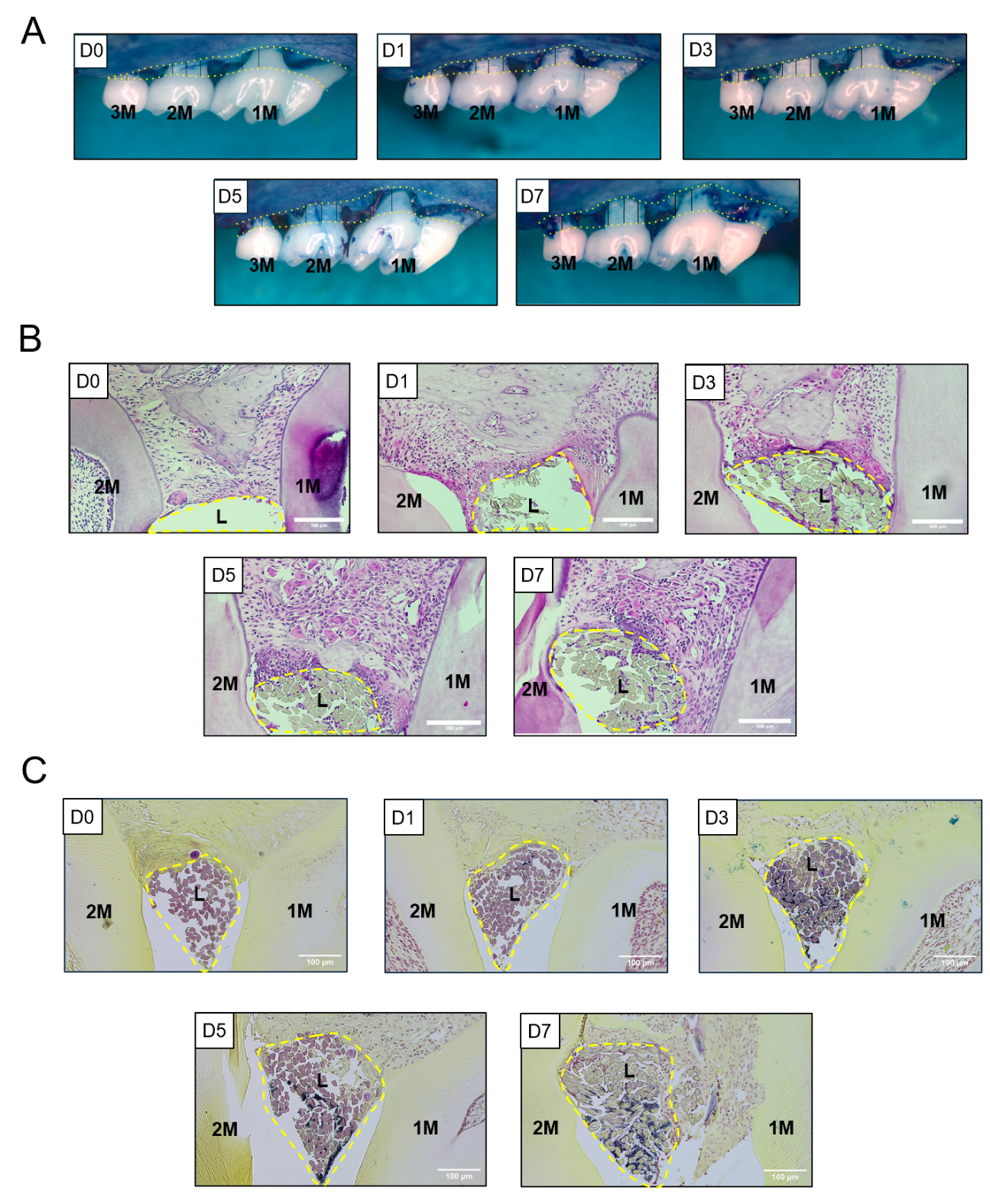

**Supplementary Figure 1.** (A) Ligature-induced periodontitis model: Images demonstrating bone measurements on ligatures at day 0, 1, 3, 5 and 7 post-induction, 6 points measured indicated by vertical lines. (B) Hematoxylin & Eosin staining show inflammatory infiltrate at day 0, 1, 3, 5 and 7 of interproximal area between first and second molars. (C) Brown & Brenn staining was performed to detect the presence of bacteria.

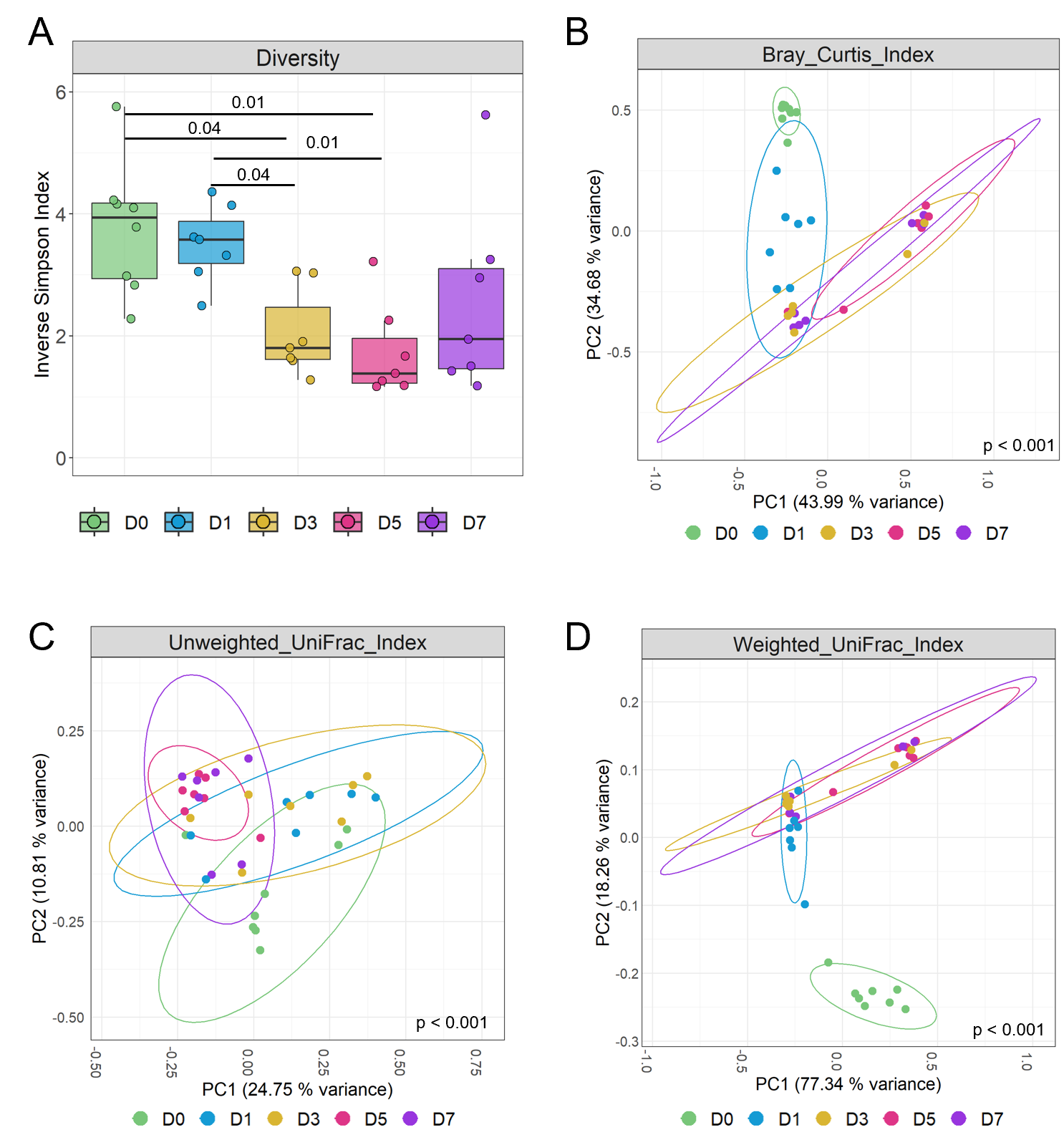

**Supplementary figure 2.** Alpha and beta diversity metrics across the timeline of ligature-induced periodontitis. (A) Inverse Simpson diversity index, a measure of alpha diversity that emphasizes the dominance of abundant taxa. Data were analysed using the Kruskal-Wallis test with Dunn's multiple comparisons post-hoc test. (B) Principal coordinate analysis (PCoA) plot constructed based on the Bray-Curtis dissimilarity index, a measure of community structure that considers abundance. Data clouds are shown with 95% confidence ellipses. The significance of the separation of data clouds was analysed using permutational analysis of variance (PERMANOVA). (C) Principal coordinate analysis (PCoA) plot constructed based on the unweighted UniFrac distance, a phylogenetic measure of community composition that considers only the presence or absence of taxa. Data clouds are shown with 95% confidence ellipses. The significance of the separation of data clouds was analysed using permutational analysis of variance (PERMANOVA). (D) Principal coordinate analysis (PCoA) plot constructed based on the Weighted UniFrac distance, a phylogenetic measure of community composition that considers the relative abundance of taxa. Data clouds are shown with 95% confidence ellipses. The significance of the separation of data clouds was analysed using permutational analysis of variance (PERMANOVA).

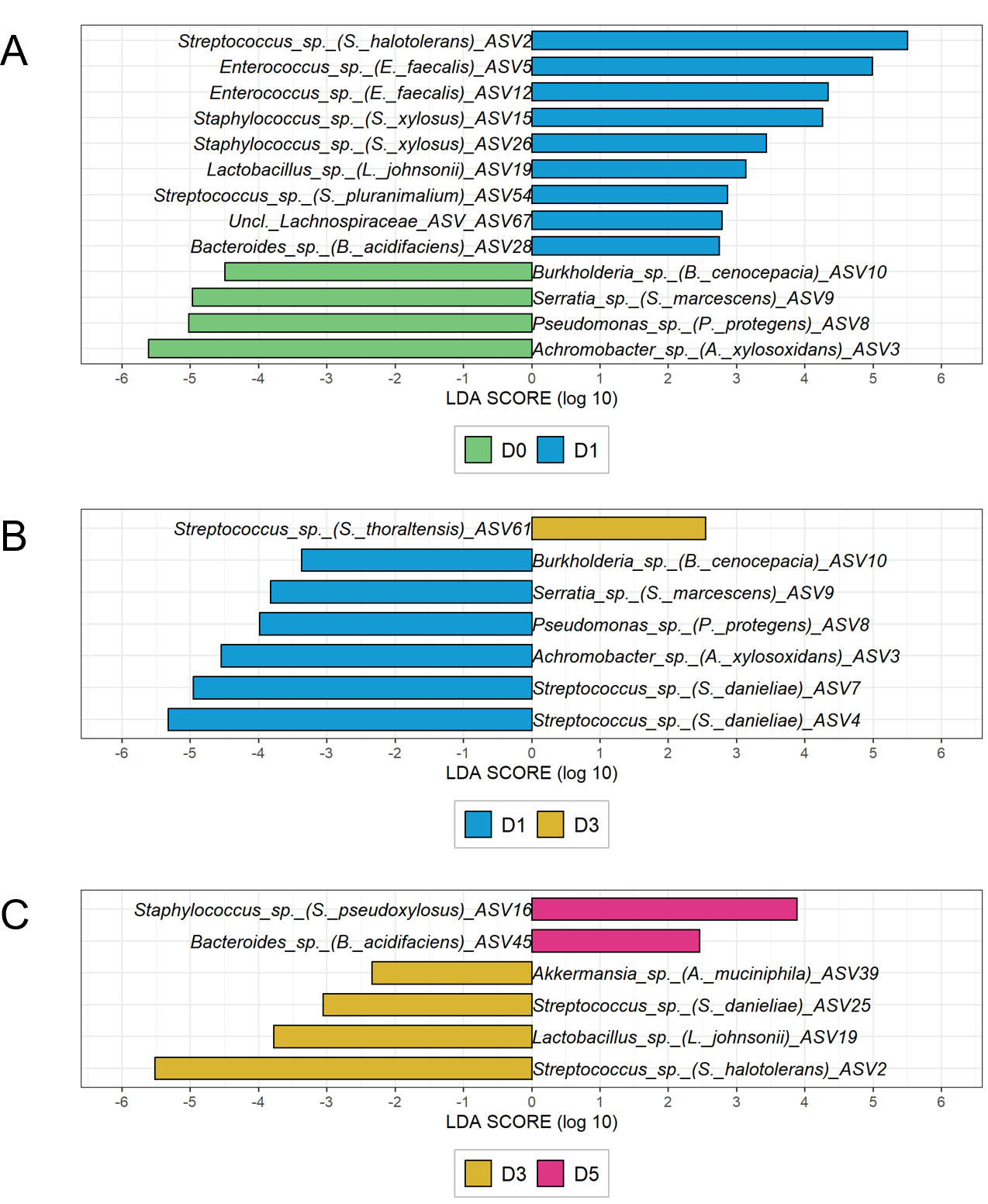

**Supplementary figure 3.** Differentially represented amplicon sequence variants (ASVs) in ligature-induced periodontitis mouse model. The graphs show taxa differentially represented according to LEfSe analyses comparing (A) D0 vs D1, (B) D1 vs D3, and (C) D3 vs D5 post-ligature. Taxa were classified at the species level. Bars represent linear discriminant analysis (LDA) scores. No differentially represented taxa were identified for the comparison D5 vs D7.

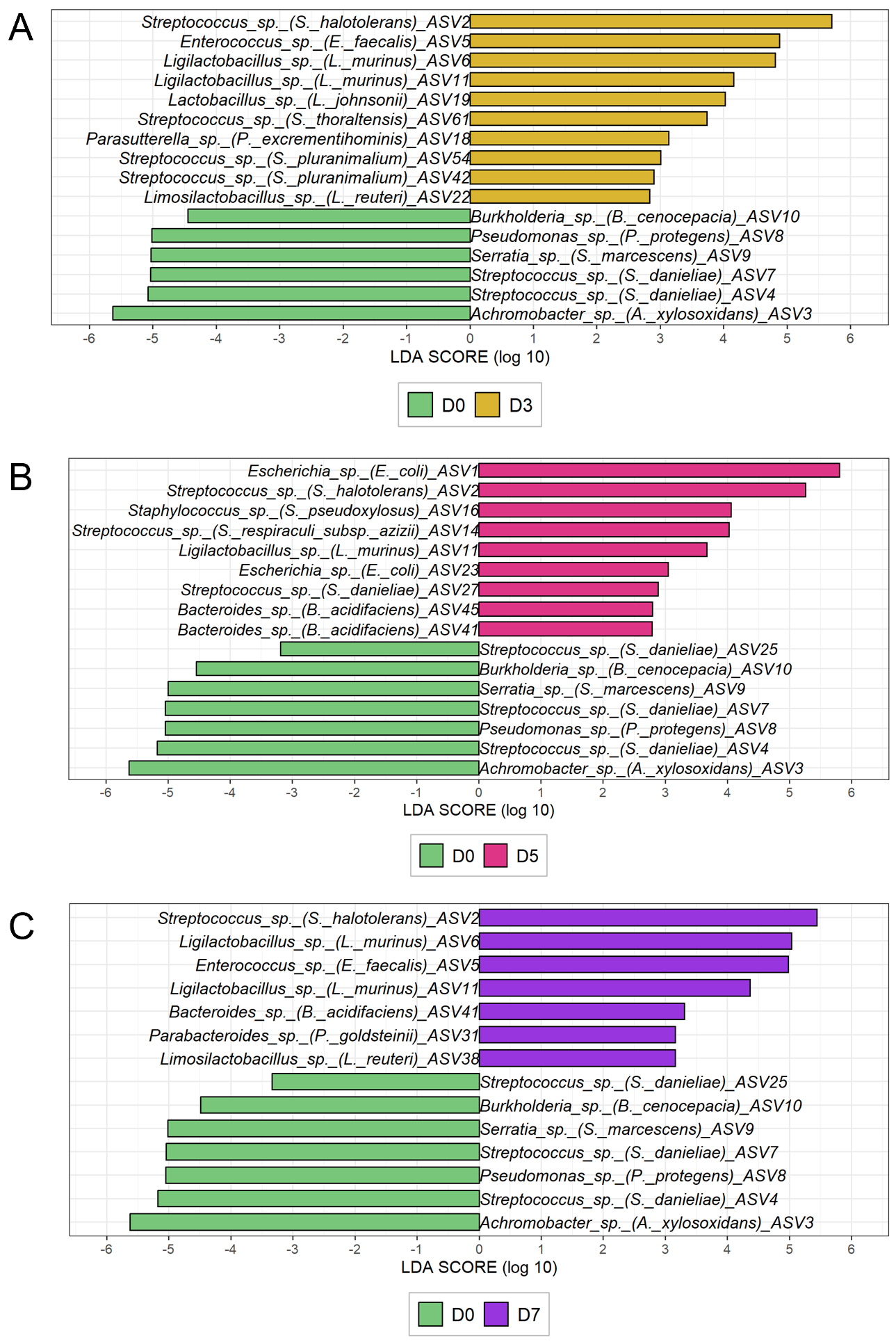

**Supplementary figure 4.**  Differentially represented amplicon sequence variants (ASVs) in ligature-induced periodontitis mouse model compared with baseline. The graphs show taxa differentially represented according to LEfSe analyses comparing (A) D0 vs D3, (B) D0 vs D5, and (C) D0 vs D7 post-ligature. Taxa were classified at the species level. Bars represent linear discriminant analysis (LDA) scores. The comparison D0 vs D1, which also showed differentially represented taxa, is presented in the Supplementary Figure 3.

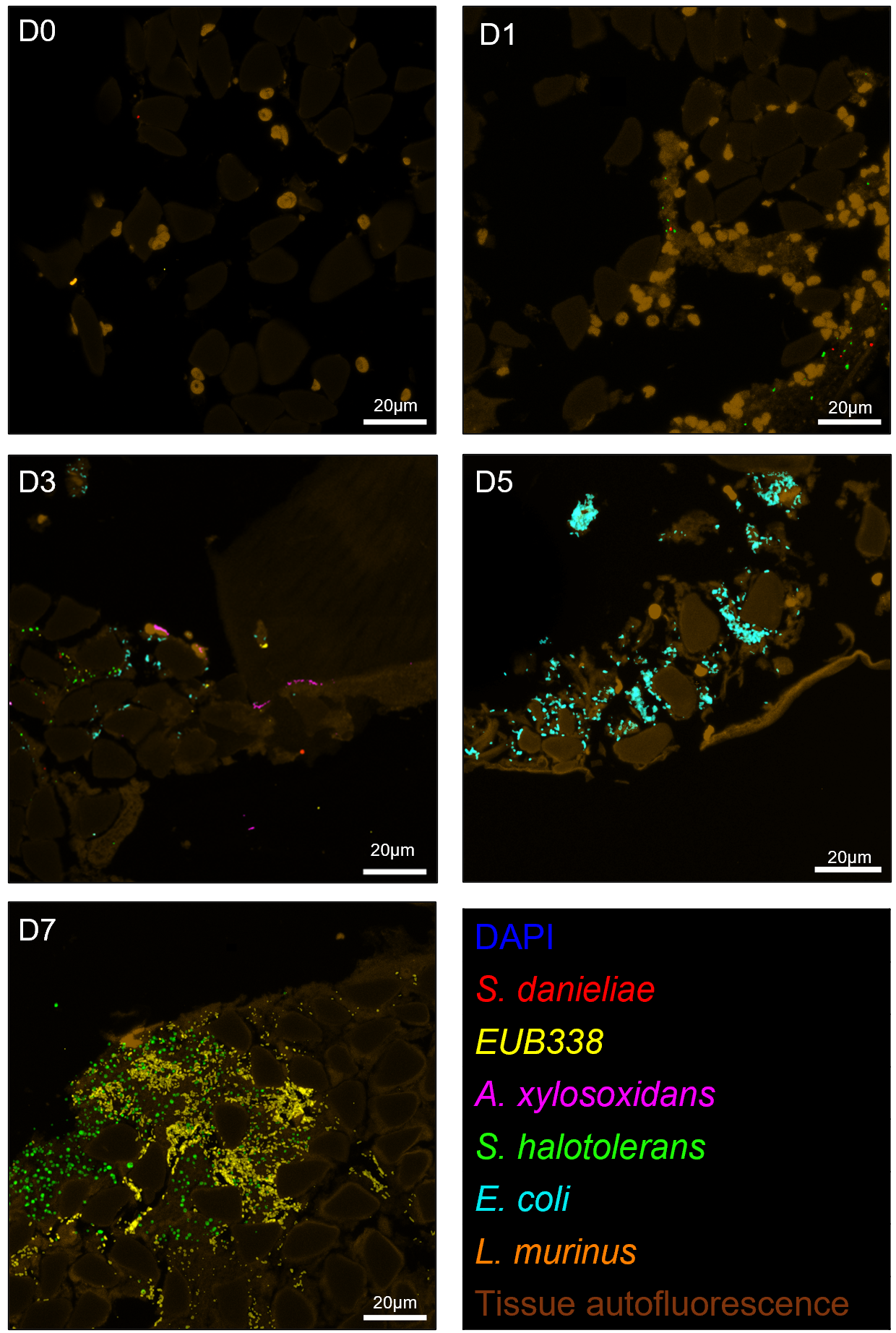

**Supplementary figure 5.** Representative CLASI-FISH images of the biofilm community on ligatures at day 0, 1, 3, 5 and 7 post-induction.

The visualized bacterial consortia correspond to the second abundant microbial profile observed in the relative abundance plot (Fig. 3B) at each respective time point. Scale bar: 20 µm. Each color represents one of the most abundant Amplicon Sequence Variants (ASVs), as identified by 16S rRNA sequencing. Probes were designed to target highly abundant ASVs.

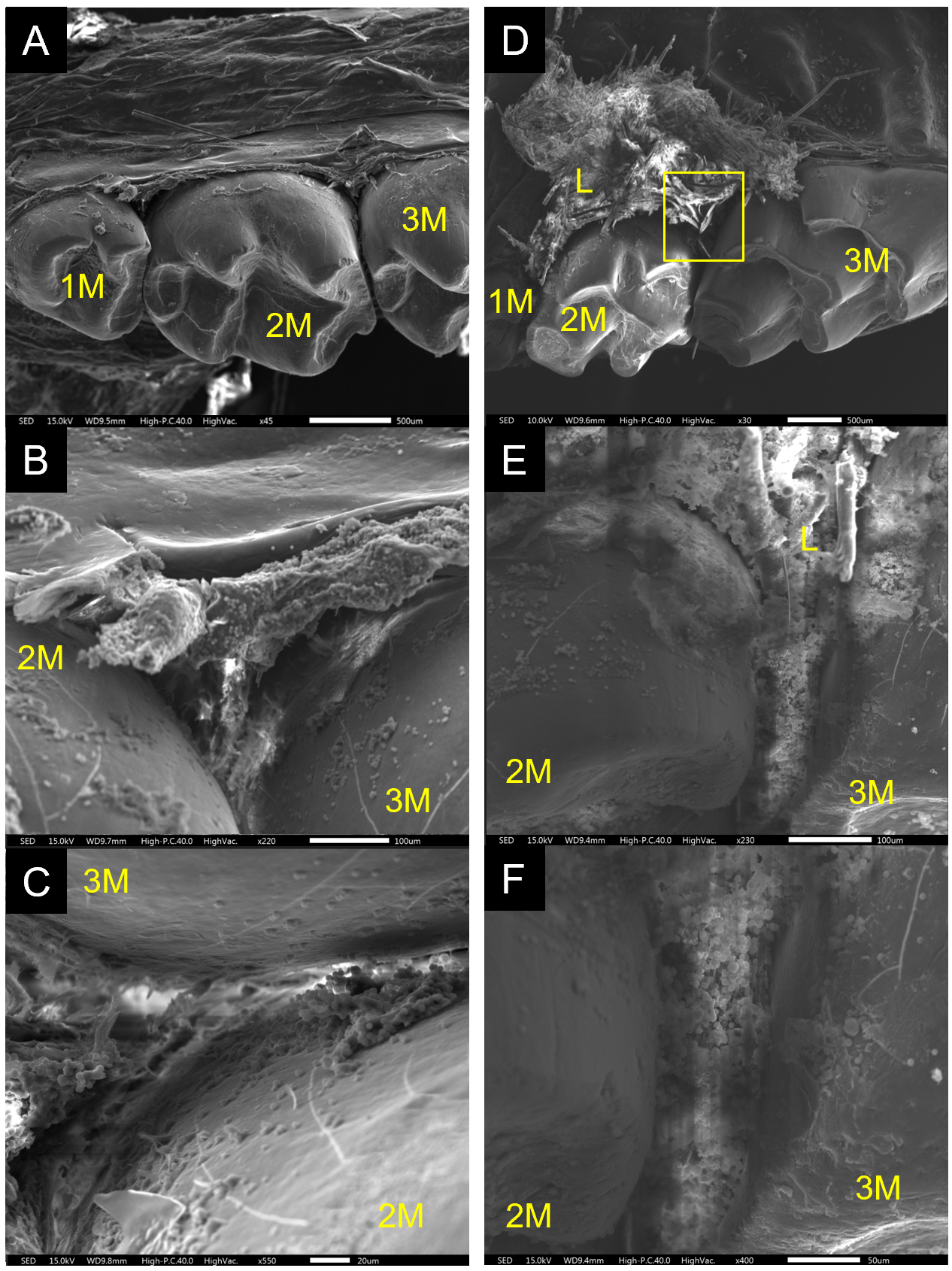

**Supplementary figure 6.** Representative scanning electron microscopy (SEM) images of the alveolar bone surface from unligated and ligature-induced periodontitis (LIP) mice. (A-C) Images of the alveolar bone surface from a non-ligated hemi-maxilla at increasing magnifications: (A) 45x, (B) 220x, and (C) 550x. (D-F) Images of the alveolar bone surface from a LIP hemi-maxilla at increasing magnifications: (D) 30x, (E) 230x, and (F) 400x. Abbreviations: M1, first maxillary molar; M2, second maxillary molar; M3, third maxillary molar; L, ligature.

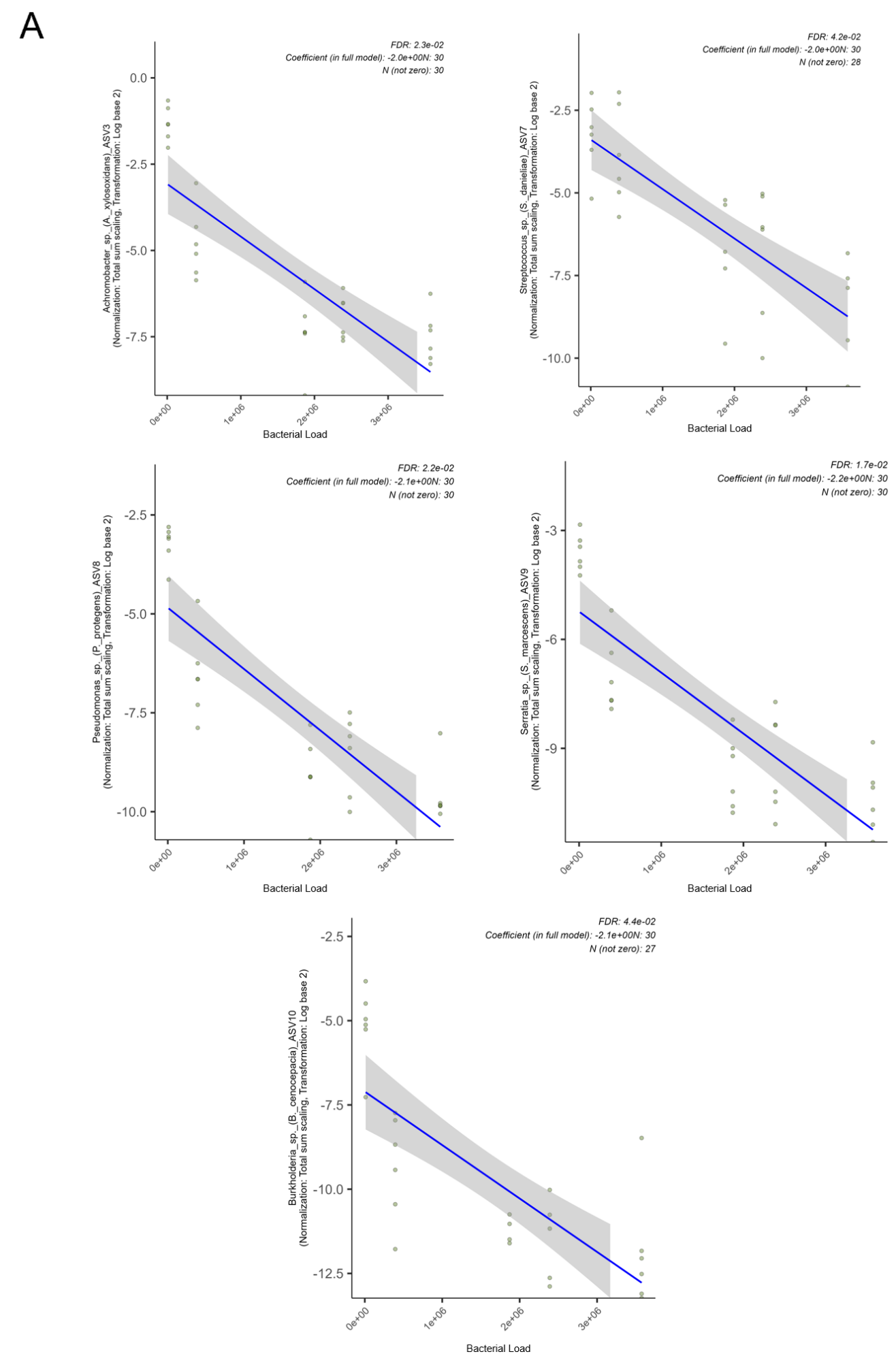

**Supplementary figure 7.** MaAsLin 3 diagnostic plot for the significant ASVs’ association with bacterial load. Scatter plot showing normalized abundance (total-sum scaling, log2-transformed) versus bacterial load. Each point represents a sample. The blue line and grey ribbon indicate the linear model fit with 95% confidence interval. Top annotations show sample size (N), number of non-zero samples, and fitted coefficient.

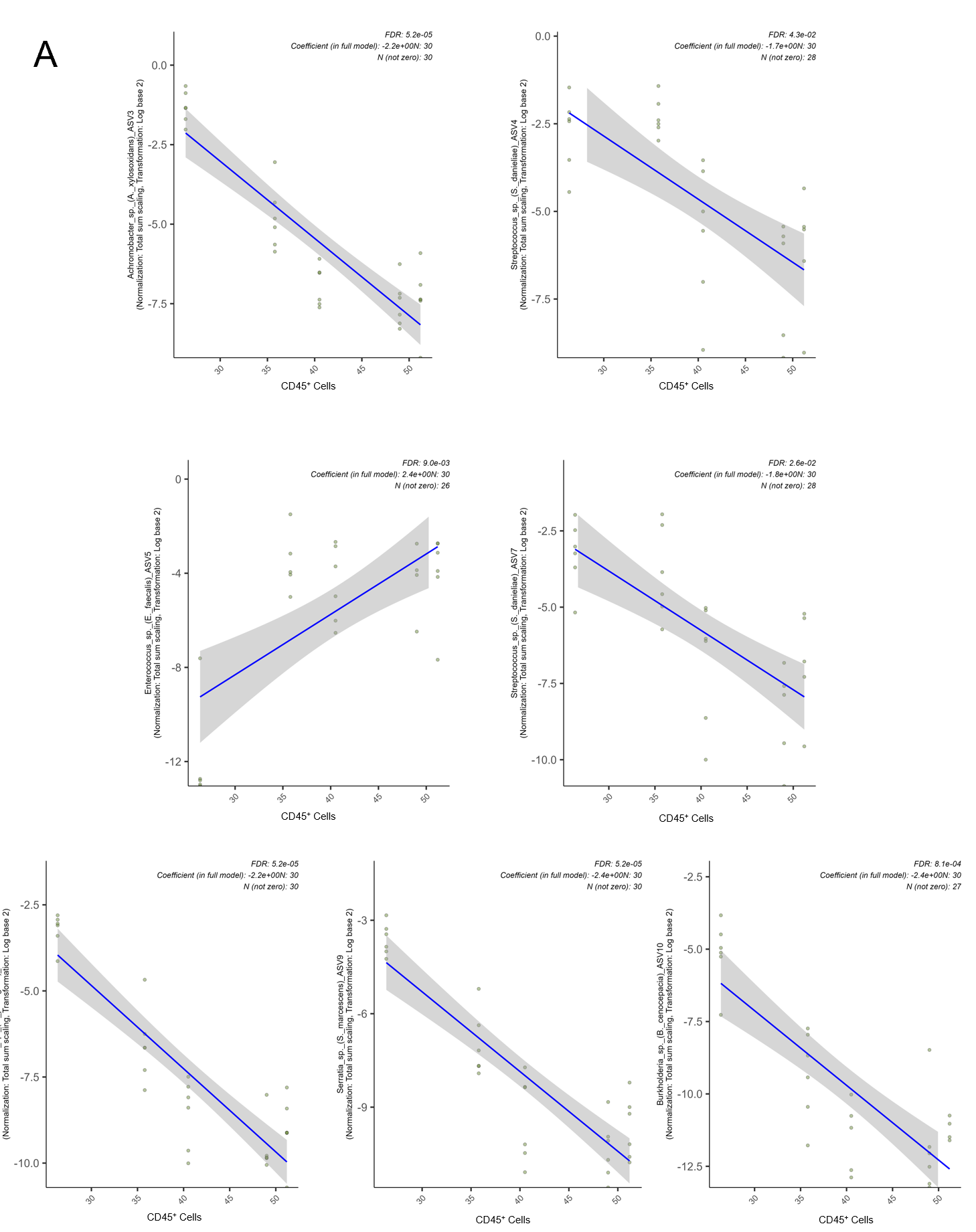

**Supplementary figure 8.** MaAsLin 3 diagnostic plot for the significant ASVs’ association with CD45+ cells. Scatter plot showing normalized abundance (total-sum scaling, log2-transformed) versus bacterial load. Each point represents a sample. The blue line and grey ribbon indicate the linear model fit with 95% confidence interval. Top annotations show sample size (N), number of non-zero samples, and fitted coefficient.

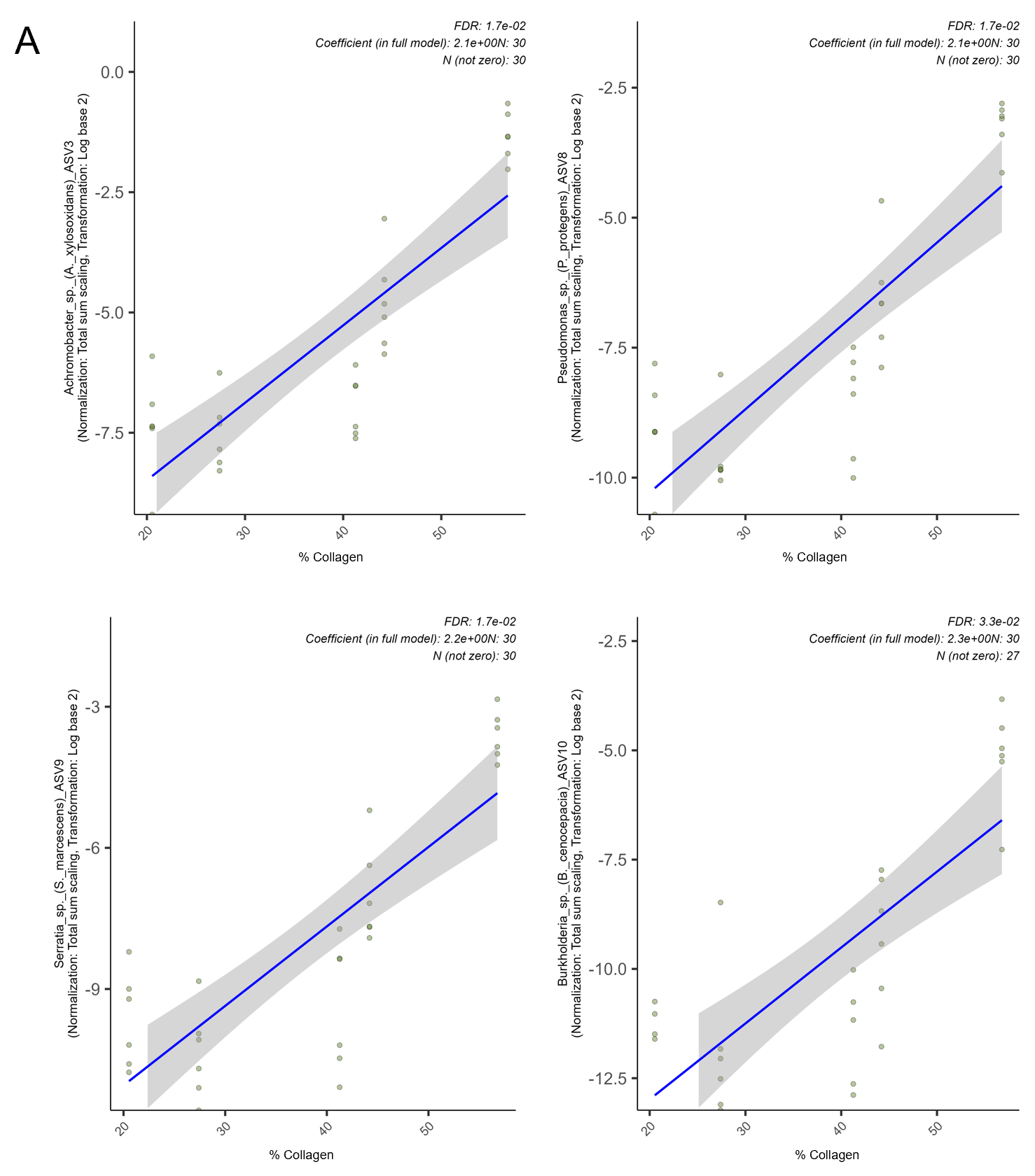

**Supplementary figure 9.** MaAsLin 3 diagnostic plot for the significant ASVs’ association with collagen percentage. Scatter plot showing normalized abundance (total-sum scaling, log2-transformed) versus bacterial load. Each point represents a sample. The blue line and grey ribbon indicate the linear model fit with 95% confidence interval. Top annotations show sample size (N), number of non-zero samples, and fitted coefficient.

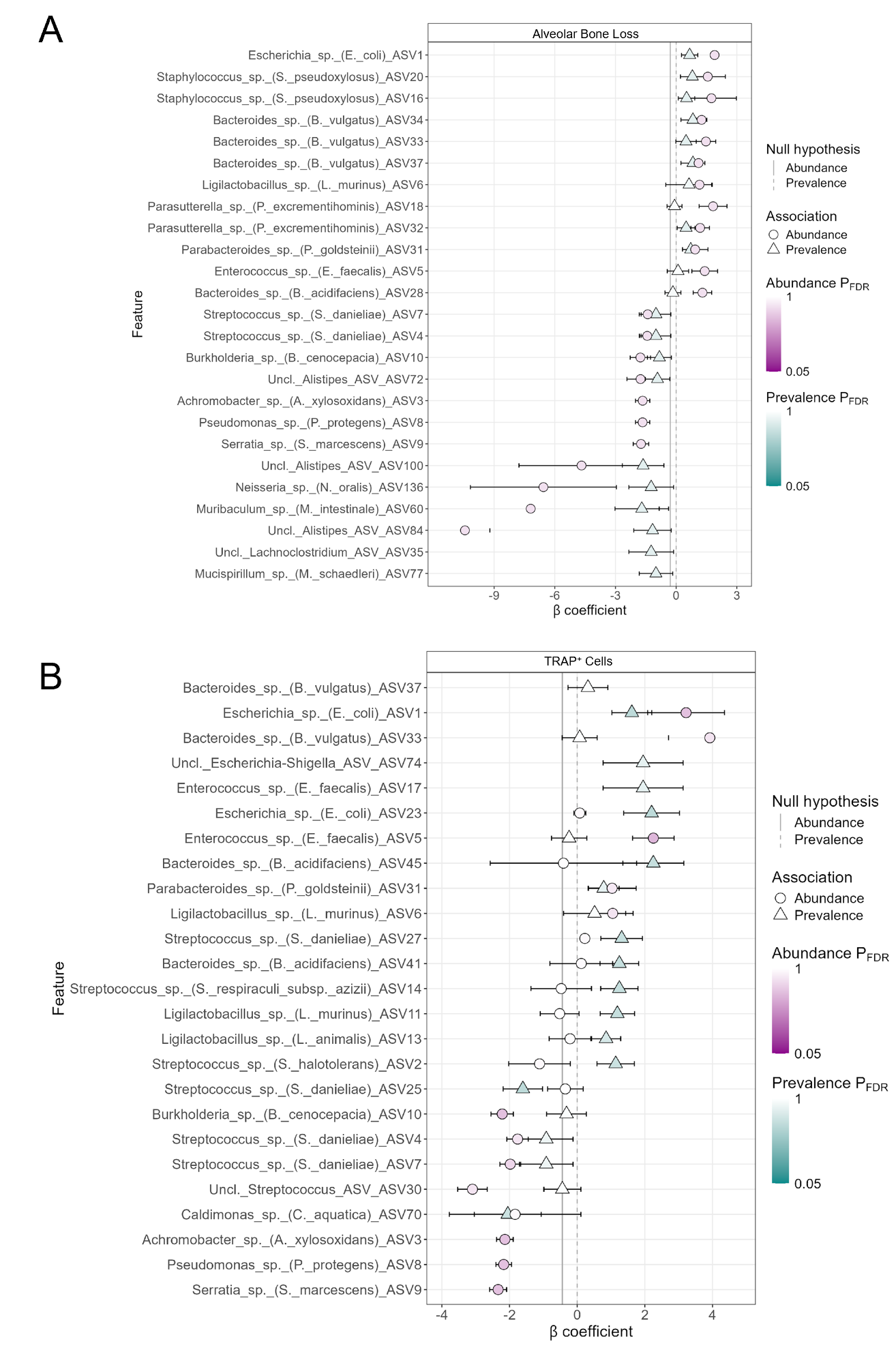

**Supplementary figure 10.** Forest plot of top microbial features associated with different variables. A. Forest plot of top microbial features associated with alveolar bone loss. B. Forest plot of top microbial features associated with TRAP+ Cells. Points represent effect sizes (coefficients) from MaAsLin 3 models, with horizontal lines indicating 95% confidence intervals. Features are ordered by significance (top = most significant). Colors distinguish abundance (blue) versus prevalence (red) associations. Negative coefficients indicate ASVs that decreased with the corresponding variable; positive coefficients indicate ASVs that increased with the corresponding variable. Labels depict ASV names.
